# Alphavirus replicons encoding IFN-γ enhance cancer virotherapy by overcoming macrophage-mediated suppression

**DOI:** 10.1101/2025.02.12.637841

**Authors:** Laura Horvathova, Priscilla Kinderman, Thijs Janzen, Baukje Nynke Hoogeboom, Franz J. Weissing, Nadine van Montfoort, Toos Daemen, Darshak K. Bhatt

## Abstract

Interference by tumor-associated macrophages may significantly reduce the efficacy of therapeutic viruses designed to infect cancer cells and activate antitumor T-cells. Using a computational model, we hypothesized that viruses encoding a T cell-stimulating signal, like IFN-γ, could overcome this barrier. We engineered an alphavirus-based replicon expressing IFN-γ and evaluated its effect in various human-derived tumor-immune coculture systems and an *in vivo* murine model. While alphavirus replicons do not replicate in macrophages, macrophages acted as a barrier, limiting tumor infection in a frequency-dependent but phenotype-independent manner. Nonetheless, T-cell activation occurred even when only a fraction of infected tumor cells expressed IFN-γ, regardless of macrophage presence, frequency, or phenotype. Additionally, viral stimulation drove macrophage-repolarization towards a pro-inflammatory phenotype favoring T-cell activation. These findings highlight a strategy for optimizing virotherapy in macrophage-rich tumors by designing viruses that stimulate T-cell activation, ensuring therapeutic efficacy.

## Introduction

Tumor-associated macrophages are a predominant component of the tumor immune infiltrate across many cancers^1^. These macrophages exhibit context-dependent phenotypic plasticity^2,3^ and function to either support or suppress antitumor immunity^2–5^. In particular, they perform phagocytosis, act as antigen-presenting cells, express checkpoint regulators, and secrete an array of cytokines that shape cancer progression and response to immunotherapy^4,6,7^.

Cancer virotherapy is a form of immunotherapy where viruses are used to infect cancer cells, and subsequently boost anticancer T cell responses^8^. Virus-induced cancer cell death causes the release of danger signals and antigens in the tumor microenvironment^9^. Ideally, these signals are immunogenic, enhancing the recruitment and proinflammatory activation of immune cells into the tumor. However, tumor-associated macrophages may hinder the therapeutic efficacy of virotherapy by capturing virus particles, thereby significantly reducing the availability of the virus for infecting cancer cells^10–13^. This effect could be particularly pronounced when macrophages are non-permissive for viral replication, as it restricts tumor cell infection and the subsequent release of immunostimulatory signals and tumor antigens, thereby indirectly suppressing downstream anticancer T-cell activity. Tumor-associated macrophages further limit oncolysis through innate antiviral signaling^10–13^ and T-cell suppression by regulatory mechanisms^3,14^. Variation in tumor composition^15,16^, particularly in terms of macrophage frequency and phenotype can lead to variability in therapeutic responses^17–19^. Enhancing T-cell-mediated antitumor responses using virotherapy thus remains challenging, especially in macrophage-rich tumors. Therefore, development of therapeutic viruses capable of strongly stimulating T-cell immune responses is crucial to overcome the limitations posed by non-permissive macrophages.

Semliki Forest virus (SFV)-based replicons are emerging as promising candidates for immunogenic virotherapy. SFV is a positive-stranded RNA virus, belonging to the *Alphavirus* genus. The RNA genome of SFV functions as a replicon, encoding non-structural viral proteins capable of viral-RNA translation and replication. Previous efforts in developing recombinant SFV particles (rSFV) have focused on improving safety by deleting viral genes that code for structural proteins^20^. This creates suicidal virus particles capable of a single round of infection. We and others have demonstrated that rSFV particles can successfully infect and express viral genes in a wide range of cancer cells and healthy stromal cells, however, macrophages are non-permissive to SFV infections and virus-encoded protein translation^21–23^. Furthermore, the immunogenicity of rSFV has been improved by encoding antigens^24–27^, cytokines^22,28^, or antibodies^29^ to boost humoral and cellular immune responses in the context of cancer therapy and vaccination against infectious diseases. Our group has demonstrated phase-1 and phase 2 clinical safety, immunogenic potential and clinical efficacy of an rSFV-based therapeutic vaccine encoding antigens of human papillomavirus in patients with cervical cancer^30,31^. So far, studies using rSFV as an anticancer agent have only been conducted in murine models^22,28,29^. Recently our group showed that rSFV can be engineered to express immunogenic human cytokines or chemokines with an enhanced potential to recruit and activate T-cells in different human cancer models^23^.

In this study, we implemented a combined theoretical and experimental approach to design an immunogenic-rSFV therapy capable of T-cell activation and macrophage polarization. First, we evaluated whether indeed macrophages limit rSFV infection in cancer cells growing in monolayer (2D, two-dimensional) or spheroid-based (3D, three-dimensional) tumor-macrophage cocultures. Second, we employed a computational model to assess whether rSFV-encoding immunogenic signals can promote anticancer T-cell responses and tumor eradication despite the presence of macrophages. Third, we analyzed literature to identify interferon-gamma (IFN-γ) as a pro-inflammatory cytokine capable of both macrophage and T-cell activation. We further evaluated the correlation between intra-tumoral IFN-γ signature and activation of macrophages and T-cells using data of cervical and pancreatic cancer patients available from The Cancer Genome Atlas (TCGA)^1^. For experimental validation, we engineered rSFV to express IFN-γ upon infection of cancer cells and employed both monolayer and spheroid-based tumor-immune cocultures to evaluate T-cell activation. In these tumor-immune cocultures, we introduced macrophages of either a naïve (M_naive_) or an IL-4-induced phenotype (M_IL4_). Finally in a proof-of-concept study, we tested the efficacy of rSFV in a pancreatic cancer mouse model. These models allowed us to assess how tumor infection and T-cell activation are influenced by macrophages and to evaluate the immunogenic potential of virus-encoded IFN-γ.

## Results

### Effect of macrophage frequency and phenotype on rSFV-mediated tumor infection

As a proof-of-concept, we performed all analyses in the context of two independent solid tumor types, i.e. cervical and pancreatic cancer, with innate differences in immune signaling (**Supplementary Figure 1**). rSFV-mediated infection of cancer cells leads to expression of encoded transgenes but not production of progeny virus particles (as illustrated in **Figure 1A**). To validate our findings in an experimental setup, we employed an *in vitro* cancer-macrophage coculture model in either a monolayer or spheroid-based spatial organization (**Figure 1B**). This allowed us to have control over factors such as the frequency and phenotype of macrophages and cancer cells. Using real-time microscopy-based imaging, we studied the effect of macrophages in regulating rSFV infection of a pancreatic cancer cell line (PANC-1). Figure 1C-D shows microscopy images resulting from the spheroid (**Figure 1C**) or monolayer (**Figure 1D**) coculture setup consisting of cancer cells and a varying frequency of macrophages. With an increasing frequency of macrophages (stained red in the images) present in the coculture, we observed a decrease in virus infection (stained green, expressing virus-encoded green fluorescent protein, GFP). We performed the cancer-macrophage coculture in various setups, either varying only the frequency of macrophages (green bar) or the frequencies of both macrophages and cancer cells (purple bar) (**Figure 1E**). In particular, we confirmed^21,32^ that virus-encoded GFP expression is restricted to cancer cells but not macrophages. The decrease in the number of infected cells corresponds to a decrease in the absolute number of target cancer cells and to a relative increase in the number of macrophages (**Figure 1F-G**), indicating that presence of macrophages limits the effect of virotherapy.

**Figure 1:**
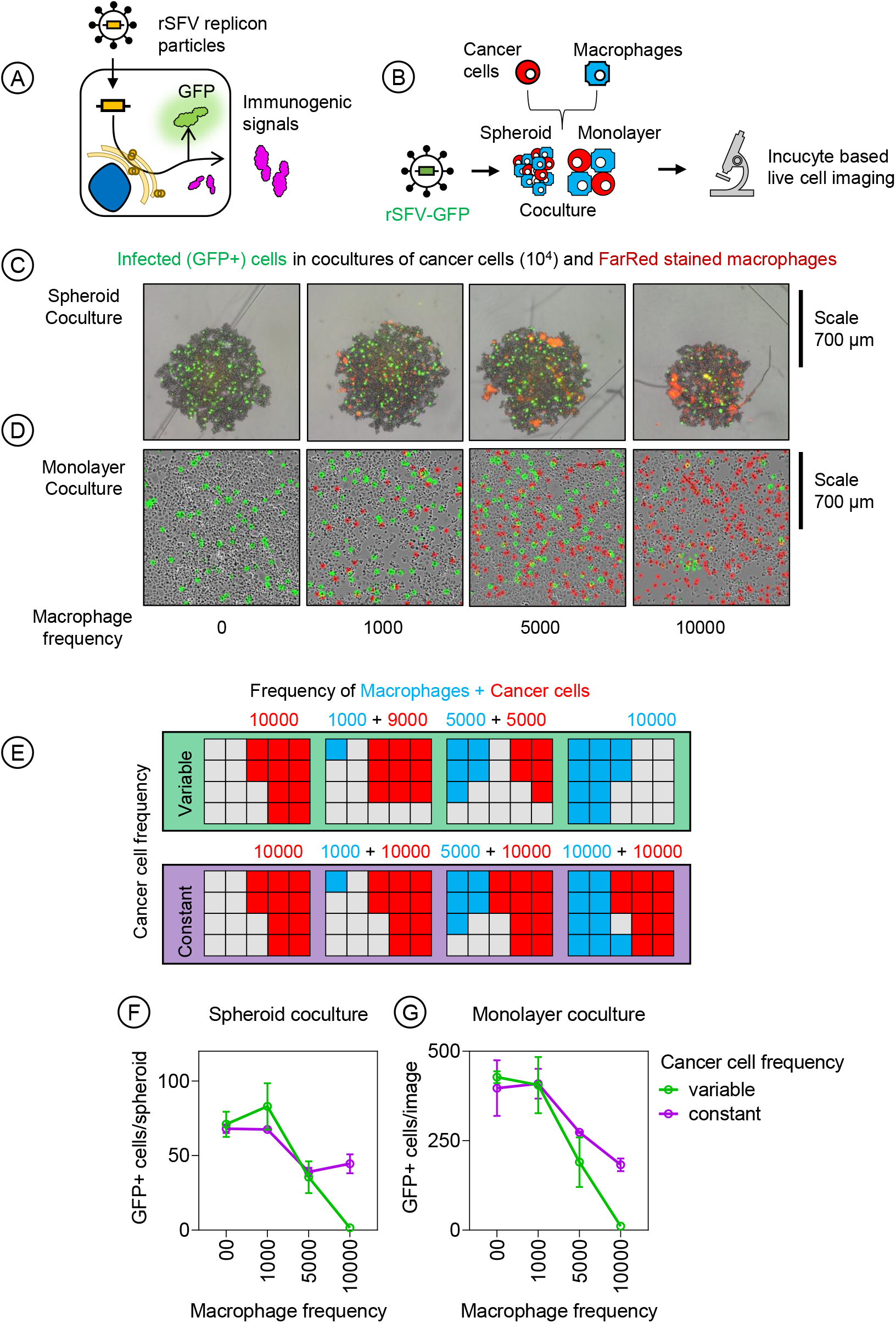
Effect of macrophage frequency on rSFV-mediated tumor infection. **(A)** Illustration explaining the mode of action of rSFV encoding either GFP or an immunogenic signal used as a non-replicating “suicidal” virotherapy. **(B)** Schematic of the experimental setup for studying rSFV infection (MOI-10) in a pancreatic cancer cell line (PANC-1) cocultured with macrophages in both monolayer (2D) and spheroid (3D) spatial organizations. The frequency of macrophages and cancer cells were controlled at the time of initiating the coculture. The coculture setup showing representative images of **(C)** spheroids or **(D)** monolayers composed of cancer cells, infected cells (green, GFP+ cells), and varying frequencies of macrophages (red, stained with FarRed dye) after 24 hours post infection with rSFV encoding GFP (rSFV-GFP). **(E)** Diagram illustrating experimental setups with varying frequencies of macrophages and cancer cells. Top row (variable setup): Frequency of macrophages and cancer cells are varied. Bottom row (constant setup): Frequency of macrophages is varied while cancer cells are kept constant. The grey cells indicate empty space that can be occupied by either cancer cells or macrophages. Quantitative analysis of GFP expression indicating virus infection in the **(F)** spheroid and **(G)** monolayer coculture setups. The plots represent data from 4 replicates of respective coculture methods. Data are presented as mean values⍰±⍰SD.

We used non-polarized macrophages for the experiments presented in Figure 1. Since tumor-associated macrophages can exhibit diverse phenotypic states, we specifically addressed the contribution of two phenotypes most frequently found in the tumor microenvironment. For this, we polarized them to either a classically activated anti-tumoral phenotype (M_class_ or M1-like) or an alternatively activated pro-tumoral phenotype (M_alter_ or M2-like) through cytokine stimulation (**Figure 2A**). Here, we used either a combination of LPS and IFN-γ to generate classically activated macrophages (M_LPS+IFNγ_), or IL-4 to generate alternatively activated macrophages (M_IL4_). Pro-inflammatory cell surface marker proteins such as CD80 and CD86 are upregulated in classically activated M_LPS+IFNγ_ macrophages, whereas regulatory markers like CD206 are more abundant in alternatively activated M_IL4_ macrophages (**Figure 2B**) also corresponding to a distinct cellular state (**Figure 2C**). We then assessed if the macrophage phenotype, in particular the alternatively activated M_IL4_ phenotype that is most frequently found in tumors, influenced tumor infection by rSFV in monolayer or spheroid cocultures (**Figure 2D-E**). We observed that the number of GFP+ cells reduced with an increase in the number of macrophages, however, independent of the macrophage phenotype (**Figure 2F-G**), as there were no differences observed in the magnitude of reduction in GFP+ cancer cells between conditions that had either M_Naïve_ or M_IL4_ polarized macrophages. This was also the case when macrophages were cocultured with a cervical cancer cell line (Ca-Ski) (**Figure 2H**).

**Figure 2:**
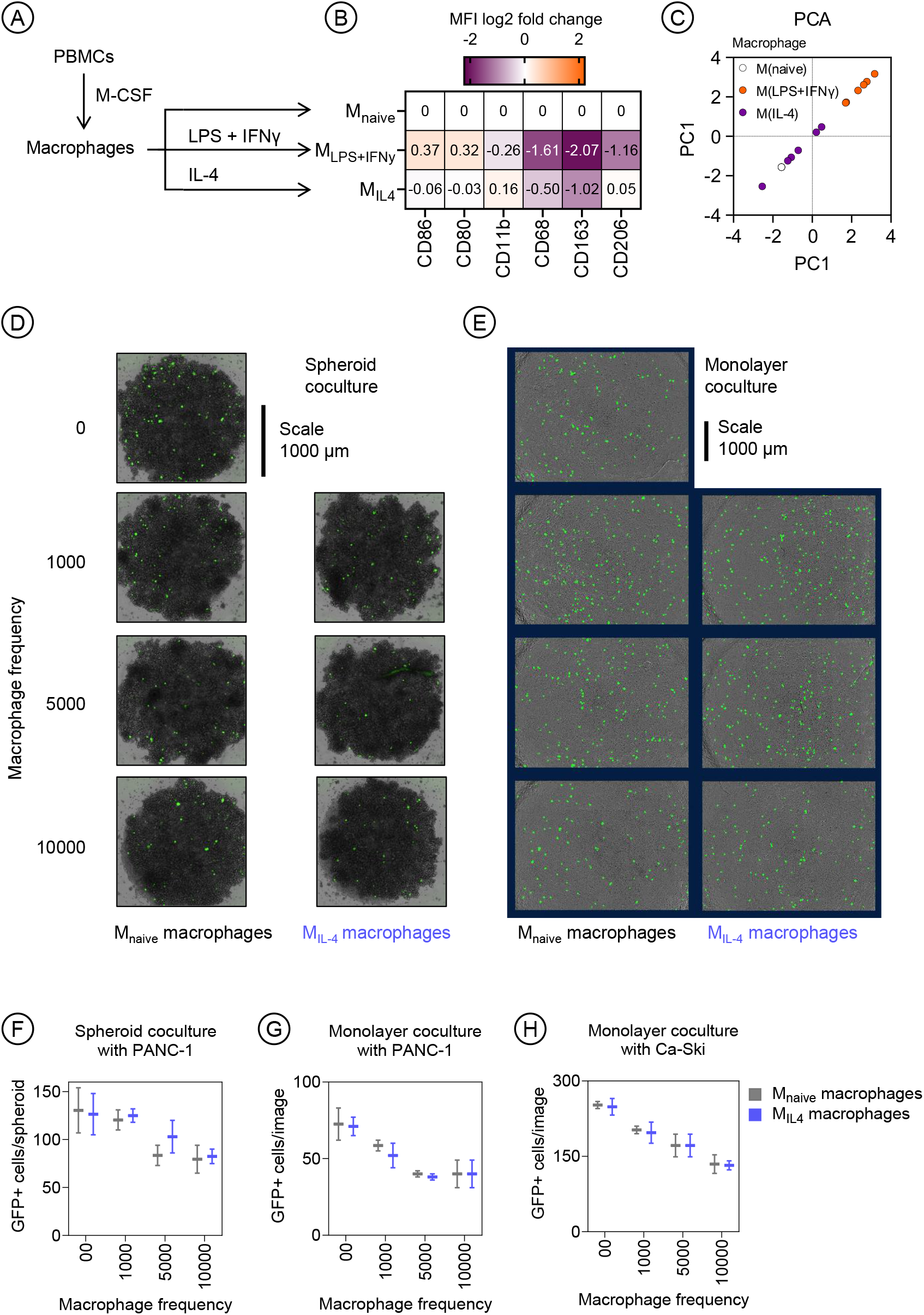
Effect of macrophage phenotype on rSFV-mediated tumor infection. **(A)** Schematic of the experimental setup for polarizing peripheral blood derived macrophages to either a naïve (M_naive_) or or an anti-tumoral (M_LPS+IFNy_ or M1-like), or a pro-tumoral (M_IL4_ or M2-like) phenotype using cytokines. **(B)** The surface expression of immunomodulatory proteins involved in anti-tumoral (CD80, CD86) and pro-tumoral (CD206, CD163) macrophage activity. **(C)** Principle component analysis of the differences in the phenotypes of macrophages from various healthy blood donors (n=5). Representative images of rSFV infection (green, GFP+ cells) in a pancreatic cancer cell line (PANC-1) cocultured with M_naive_ and M_IL4_ macrophages in both spheroid **(D)** and monolayer **(E)** spatial organizations. Quantitative analysis of GFP expression indicating virus infection (MOI-10) in the **(F)** spheroid and **(G)** monolayer coculture of PANC-1 cells with M_naive_ and M_IL4_ macrophages. **(H)** Quantification of GFP expression in monolayer coculture of Ca-Ski cancer cells with M_naive_ and M_IL4_ macrophages. The plots represent data from 4 replicates of respective coculture methods. Data are presented as mean values⍰±⍰SD.

Notably, cytokine-driven macrophage polarization had no impact on the expression of genes encoding SFV-entry receptors (**Supplementary Figure 2**). Furthermore, blocking JAK-STAT signaling to examine the role of type-I interferon in antiviral effects failed to improve rSFV infection in cancer cells (**Supplementary Figure 3**). Thus, the frequency of non-permissive macrophages, rather than antiviral signaling or their polarized phenotype, primarily contributed to the observed reduction in infection.

### Model predictions regarding the effectiveness of immunogenic-rSFV therapy

In a previous study, we developed a spatiotemporal model to assess the effect of anticancer T-cell responses in response to immunogenic signals released upon virotherapy^33,34^. **Figure 3A** (left image) depicts a snapshot of a computer simulation with a tumor containing uninfected cancer cells, virus-infected cancer cells, and macrophages. Upon infection-caused cell death, the model assumes that virus-induced immunogenic signals are released in the neighborhood (as depicted in **Figure 3A** right and illustrated in **Figure 3B**) stimulating anticancer T-cell response. Using this model, we assessed the efficacy of rSFV encoding immunogenic signals as a non-replicating suicidal virotherapy. To estimate therapeutic success, we assessed the probability of tumor eradication caused by rSFV-infection and anticancer T-cell cytotoxicity. Panels *C* to *E* show, for five scenarios regarding the percentage of initially infected tumor cells (0-50%), how the probability of tumor eradication is related to the IFN-γ level produced per infected cell (**Figure 3C**), the degree of T-cell cytotoxicity (**Figure 3D**), and the frequency of macrophages at the time of rSFV therapy (**Figure 3E**). Generally, 30% initially infected cells are sufficient to promote tumor eradication even at low levels of immunogenic signal production. A minimum T-cell cytotoxicity level of 3 (corresponding to 3 target cells killed per cytotoxic T-cell per day) is required for an effective therapeutic outcome, as increasing numbers do not correspond with increased tumor eradication. Therapeutic outcome was found to be mainly determined by the percentage of initially infected cells producing immunogenic signals and not by the macrophage frequency. To guide the selection of effective immunogenic signals for rSFV-based therapy, we next consulted the literature to identify candidates with strong T-cell-activating potential.

**Figure 3:**
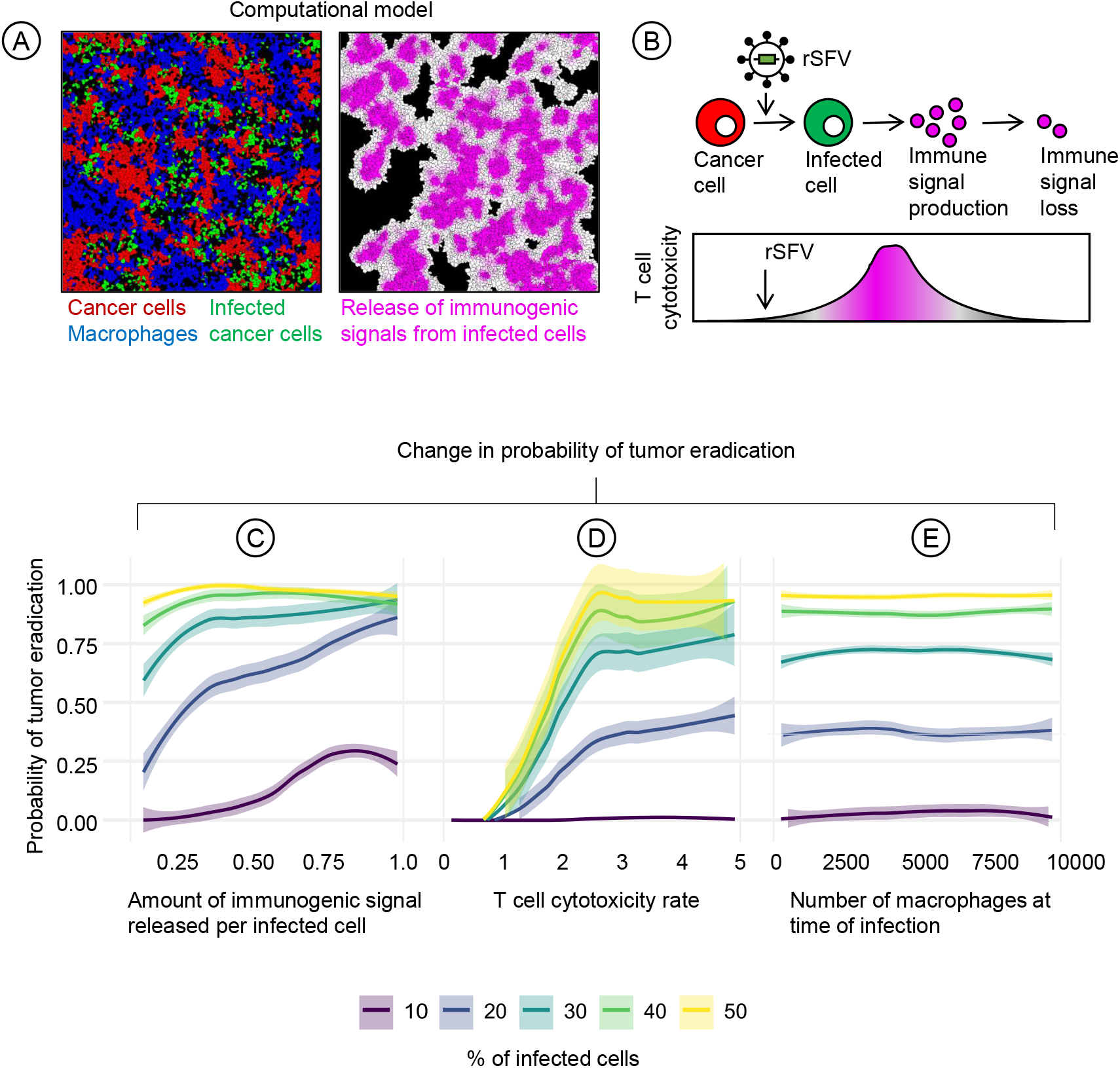
Predicting the outcomes of immunogenic-rSFV therapy using a spatiotemporal model of tumor development. **(A)** Snapshot of a simulation. Left: Spatial structure of a tumor containing uninfected cancer cells (red), virotherapy-infected cancer cells (green), and macrophages (blue). Right: Distribution of infection-induced immunogenic signals (magenta) released from dying infected cancer cells, which stimulate a cytotoxic anti-cancer T-cell response. **(B)** Time trajectory illustrating the dynamics of a suicidal rSFV-therapy. Initially, uninfected cancer cells are targeted by the virus and weakly attacked by T-cells. Over time, virus-killed cancer cells release immunogenic signals, boosting T-cell cytotoxicity. Finally, the boosting effect wanes as immunogenic signal-levels diminish. **(C)** Effect of the amount of immunogenic signal produced per infected cells on the probability of tumor eradication. The colors indicate five simulation scenarios differing in the percentage of infected cells at the time of infection. **(D)** The effect of anticancer T-cell cytotoxicity rates on the probability of tumor eradication. **(E)** The effect of macrophage frequency at the time of rSFV therapy on the probability of tumor eradication. When not specifically under consideration, the parameter values were kept at their default values: the amount of immunogenic signal released was set to 0.25, the T-cell cytotoxicity rate was set to 5, and the number of macrophages was set to 2500. Colored dotted lines indicate the mean values, colored envelopes indicate the 95% confidence interval obtained via bootstrapping. Each panel represents 10000 simulations for respective parameter combinations.

### The association between IFN-γ and macrophage-T-cell activation

We analyzed current literature to screen potential immunomodulatory cytokines produced and/or consumed by T-cells and macrophages. By re-evaluating previously published data^35^, we found that IFN-γ produced by T-cells represents a unique antitumoral cytokine that has the potential to stimulate both macrophages and T-cells through JAK-STAT signaling pathway (**Figure 4A**) with an effective concentration ranging from 3 to 40 pM (**Figure 4B**).

**Figure 4:**
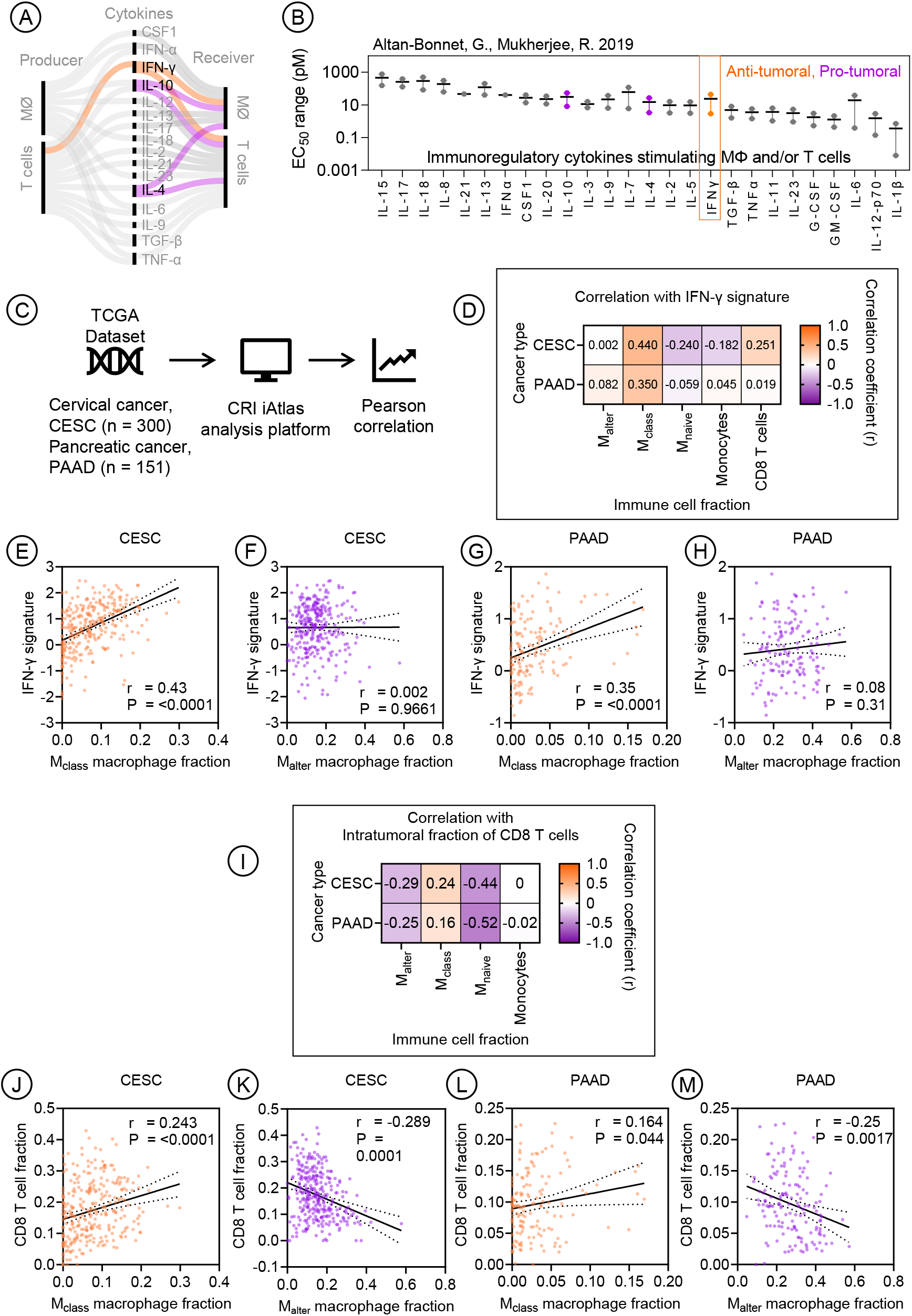
Impact of IFN-γ on tumor associated macrophage and T-cells in the tumor microenvironment. **(A)** Schematic of cytokine production and reception by T-cells and macrophages (MØ), highlighting IFN-γ, IL-10, and IL-4 as key cytokines involved in influencing both cell types. Cytokine interactions are color-coded based on their impact on tumor progression: pro-tumoral (purple) and antitumoral (orange). **(B)** The effective concentration range of various cytokines (EC_50_) with the points indicating the maximum and minimum experimentally reported limits (as per ^35^). **(C)** Analysis process of cervical (CESC, 300 samples) and pancreatic (PAAD, 151 samples) cancer patient datasets stored at The Cancer Genome Atlas (TCGA) by using the CRI iAtlas online platform to correlate the presence of macrophage types with T-cells and IFN-γ signature. **(D)** Correlation heatmap of intra-tumoral IFN-γ signature with presence of various macrophage phenotypes. **(E-H)** Scatter plots showing the relationship between the intra-tumoral IFN-γ signature with the presence of classical (M_class_ or M1-like) and alternatively activated (M_alter_ or M2-like) macrophages in cervical or pancreatic cancer patients. **(I)** Correlation heatmap of intra-tumoral fraction of cytotoxic (CD8) T-cells with presence of various macrophage phenotypes. **(J-M)** Scatter plots showing the relationship between the intra-tumoral fraction of CD8 T-cells with the presence of M_class_ or M_alter_ macrophages in cervical or pancreatic cancer patients. In E-F and J-M, bivariate linear regression was performed to assess correlation, with r (correlation coefficient) and p-values indicated per plot.

Based on cervical and pancreatic cancer patient data from TCGA^1,36^, we analyzed the correlation between the IFN-γ signature in tumor samples with the frequency of macrophages and T-cells and the phenotype of macrophages (**Figure 4C**). The IFN-γ signature applied here was derived from the TCGA pan-cancer immunogenomic analysis^1,36^. **Figure 4D** shows an overview of the correlation between the intratumoral IFN-γ signature and different phenotypes of macrophages and cytotoxic CD8 T-cells. The IFN-γ signature was found to positively correlate with a classically activated (M_class_ or M1-like) pro-inflammatory macrophage phenotype in both cervical (CESC) and pancreatic (PAAD) tumor samples (**Figure 4E, 4G**) but not with an alternatively activated (M_alter_ or M2-like) immunosuppressive macrophage phenotype (**Figure 4F, 4H**). As T-cells are one of the primary cells producing IFN-γ, we analyzed if there was a correlation between intra-tumoral CD8 T-cells and a particular macrophage phenotype (**Figure 4I**). We observed a positive correlation between classically activated (M_class_) proinflammatory macrophages and the frequency of intra-tumoral CD8 T-cells (**Figure 4J, 4L**). Inversely, there was a negative correlation between alternatively activated (M_alter_) immunosuppressive macrophages and CD8 T-cell frequency (**Figure 4K, 4M**). These findings highlight a link between IFN-γ–driven macrophage polarization and CD8 T-cell infiltration, prompting further investigation into whether rSFV-delivered IFN-γ could enhance this interaction in tumor–immune cocultures.

### T cell activation by rSFV in the presence of macrophages

We treated monolayer cocultures of macrophages and cancer cells with rSFV and added peripheral blood mononuclear cells (PBMCs) to the culture to introduce T-cells and evaluate their activation (**Figure 5A**). Through flow cytometry, we quantified expression of immune activation (CD69) and cytotoxic-degranulation (CD107a) markers as a proxy of CD4 (helper) and CD8 (cytotoxic) T-cell activation. These markers were chosen from a broader panel of T-cell activation and regulation markers (**Supplementary Figure 4**,**5**). T-cells were considered fully activated when simultaneously positive for CD38, CD69, and CD107a expression. **Figure 5B and 5C** illustrate the population distribution of activated CD4 and CD8 T-cells in different scenarios of rSFV infection and tumor-immune cocultures consisting of either PANC-1 cells or Ca-Ski cells in culture with M_naive_ or M_IL4_ macrophages in different frequencies. The upper right quadrant in each flow cytometry panel indicates the population of fully activated T-cells as characterized by the expression of CD69 and CD107a.

**Figure 5:**
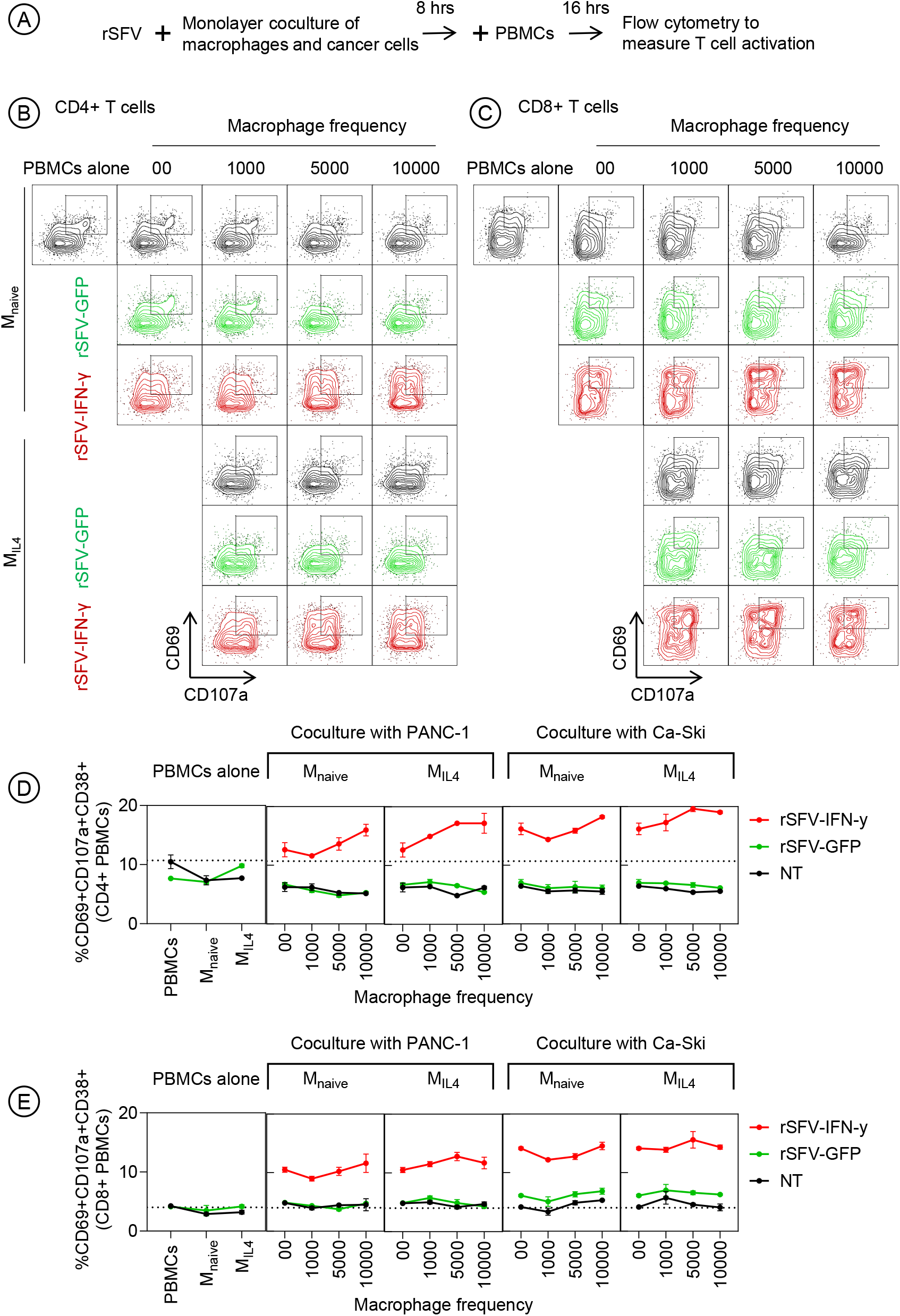
Evaluation of T-cell activation in tumor-immune monolayer cocultures by rSFV. **(A)** Schematic representation of the experimental setup consisting of tumor-immune monolayer cocultures of cancer cells and macrophages treated with rSFV (MOI-10), followed by the addition of PBMCs to introduce T-cells and evaluate their activation. Flow cytometry analysis to measure activation of **(B)** CD4 helper T-cells and **(C)** CD8 cytotoxic T-cells based on the expression of CD69 and CD107a proteins, marked in the upper right quadrant in each panel. rSFV-GFP treated cocultures are indicated in green, rSFV-IFN-γ treated cocultures in red and non-infected (NT) in black. The representative plots in (B-C) illustrate T-cell activation in the context of PANC-1 cancer cells in coculture with either M_naive_ or M_IL4_ macrophages in different frequencies. Quantification of **(D)** CD4 helper T-cell or **(E)** CD8 cytotoxic T-cell activation in the tumor-immune cocultures by rSFV. The plots represent data from 4 replicates. Data are presented as mean values⍰±⍰SD.

We observed a decrease in CD4 but not CD8 T-cell activation as a result of coculturing PBMCs solely with either M_naive_ or M_IL4_ macrophages (black circle/line, **Figure 5D**, 6E leftmost panel). PBMC coculture with cancer cells alone or in the presence of increasing frequency of either M_naive_ or M_IL4_ macrophages also led to a similar decrease in CD4 T-cell activation when compared to PBMCs alone (NT = no treatment, black circle/line, **Figure 5D**). rSFV particles themselves in the absence of target cancer cells did not have any immunomodulatory effects on CD4 or CD8 T-cell activation directly (green circle/line, **Figure 5D, 5E**, leftmost panel). Upon rSFV-GFP therapy, i.e. not encoding IFN-γ, CD8 T-cell activation was only observed in the case of infecting Ca-Ski cells but not PANC-1 cells and was independent of macrophage presence or phenotype (green circle/line, **Figure 5E**, rightmost panel). Upon rSFV-IFN-γ (rSFV-encoding IFN-γ) treatment, however, both CD4 and CD8 T-cells were activated independently of infecting Ca-Ski cells or PANC-1 cells (red circle/line, **Figure 5D, 5E**). Here, CD4 T-cell activation increased with an increasing number of either M_naive_ or M_IL4_ macrophages present in the coculture (**Figure 5D**). We confirmed that rSFV-encoded IFN-γ mediates T cell activation by inhibiting JAK-STAT signaling in T cells. Treatment with ruxolitinib, in combination with rSFV-IFN-γ, resulted in a reduced frequency of activated T cells as compared to when treated with rSFV-IFN-γ alone (**Supplementary Figure 6**).

Next, we assessed T-cell activation in spheroid-based tumor-immune cocultures (**Figure 6A**). In contrast to the results from the monolayer-based coculture system, we observed a decrease in the number of activated CD4 T-cells with an increasing frequency of macrophages in the non-rSFV-treated coculture (black circle/line, **Figure 6B**). However, similar to the results from the monolayer coculture, we observed that rSFV-GFP particles only induced CD8 T-cell activation in the case of infecting Ca-Ski cells but not PANC-1 cells (green circle/line, **Figure 6C**, rightmost panels). Moreover, rSFV-IFN-γ also led to a higher frequency of activated CD4 and CD8 T-cells independent of whether it infected Ca-Ski cells or PANC-1 cells (red circle/line, **Figure 6B, 6C**). Here, CD4 and CD8 T-cell activation even improved further with an increasing frequency of either M_naive_ or M_IL4_ macrophages.

**Figure 6:**
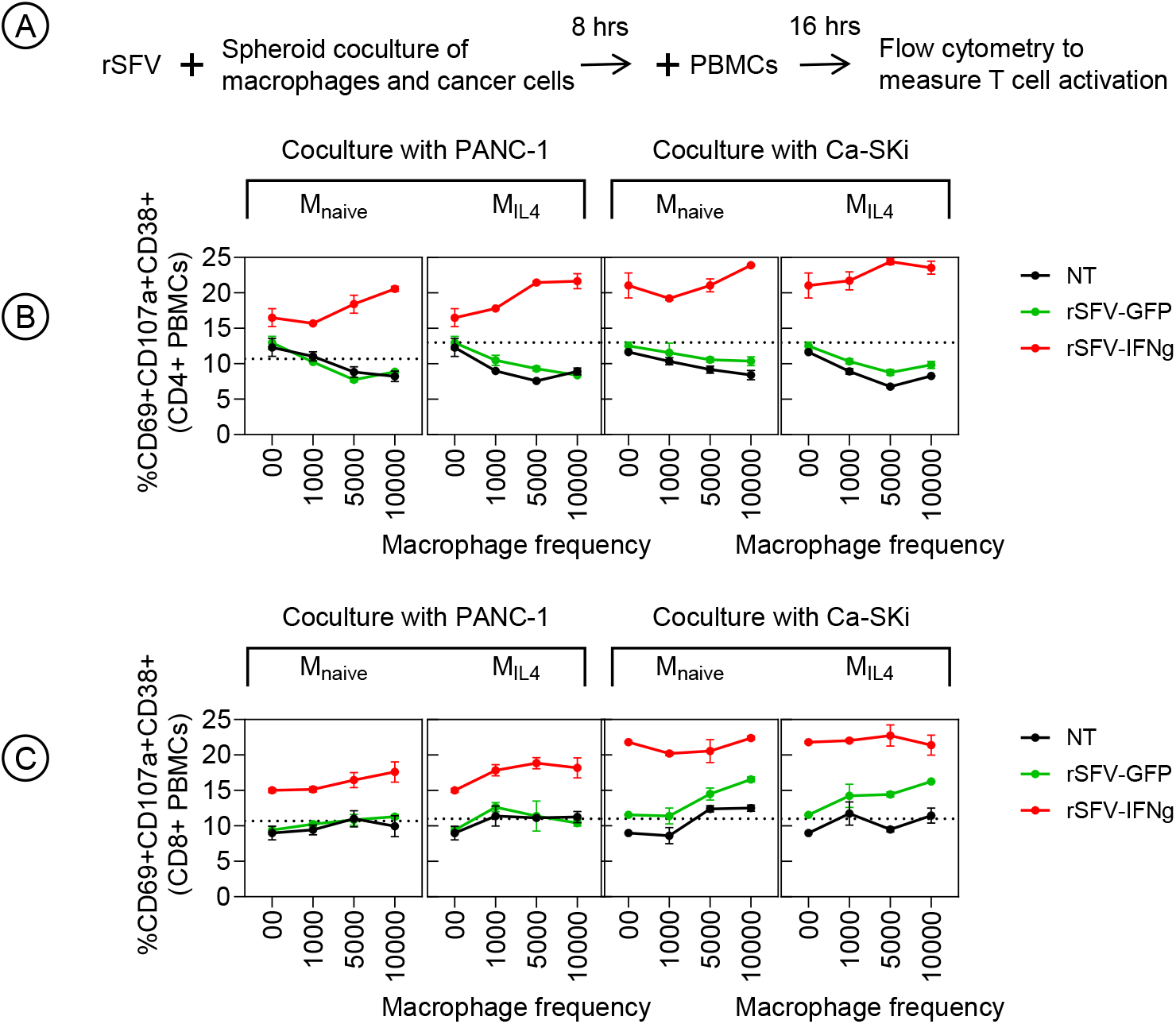
Evaluation of T-cell activation in tumor-immune spheroid co-cultures upon rSFV infection. **(A)** Schematic representation of the experimental setup consisting of tumor-immune spheroid cocultures of cancer cells and macrophages treated with rSFV, followed by the addition of PBMCs to introduce T-cells and evaluate their activation. Flow cytometry analysis to measure activation of T-cells as explained in Figure 6. Quantification of **(B)** CD4 helper T-cell or **(C)** CD8 cytotoxic T-cell activation in the tumor-immune cocultures by rSFV. rSFV-GFP treated cocultures are indicated in green and rSFV-IFN-γ treated cocultures in red. The plots represent data from 4 replicates. Data are presented as mean values⍰±⍰SD.

### Multimodal activation of macrophages by rSFV-IFN-γ

Recently our group has shown that rSFV replicon virus particles have a direct influence on macrophage phenotype and can lead to their pro-inflammatory activation, in part by JAK-STAT and NF-κB signaling, independent of their initial state^32^. Consequently, we evaluated if macrophages are also stimulated by infected cancer cells alone and if there is any additional effect in combination with virus-mediated activation (**Figure 7A**). To quantify this response, we assessed the change in expression of costimulatory molecules (CD80, CD86) on macrophages involved in T-cell regulation in addition to CD206 expression that is often associated with a regulatory phenotype and poor cancer prognosis. IL-4 polarized (M_IL4_) regulatory macrophages were found to upregulate CD86 and CD80 cell-surface expression upon stimulation with either only rSFV particles (**Figure 7B**) or only PANC-1 infected cells expressing virus-encoded IFN-γ (**Figure 7C**) or both (**Figure 7D**). The upregulation of CD80 and CD86 was the highest when macrophages received simultaneous stimulation by virus particles and infected cells producing virus-encoded IFN-γ. Importantly, significant downregulation of cell-surface CD206 expression was observed when macrophages were stimulated with infected cells producing virus-encoded IFN-γ but not GFP. Notably, IL-4 polarized macrophages upregulated the T-cell inhibitory ligand PD-L1 in the presence of cancer cells infected with rSFV-IFN-γ (**Supplementary Figure 7**). This upregulation is likely driven by canonical IFN-γ signaling pathways. However it is important to highlight that this phenomenon did not coincide with a reduction of T-cell activation phenotype as shown in **Figure 5** and **6**.

**Figure 7:**
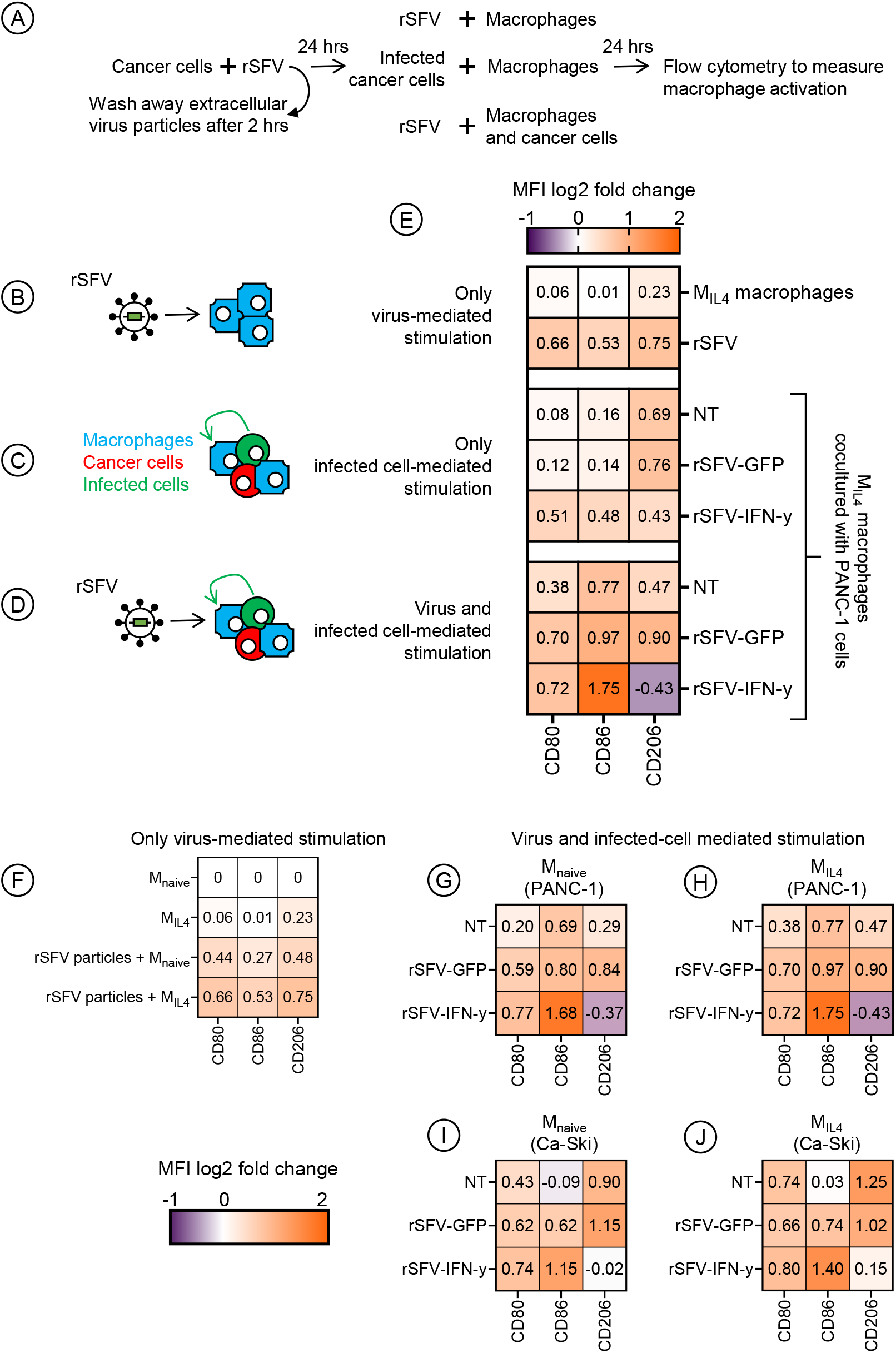
Influence of rSFV infection on macrophage phenotype. **(A)** Experimental setup to study changes in macrophage phenotype upon stimulation by rSFV (MOI-10). This includes studying the effect of **(B)** either rSFV-particles alone (rSFV row), **(C)** or by infected cells alone, **(D)** or by both. **(E)** Heatmap of change in cell-surface expression of proteins involved in either pro-tumoral (CD206) or anti-tumoral (CD80, CD86) function of M_IL4_ macrophages cocultured or not with PANC-1 cancer cells upon stimulation resulting from rSFV particles and/or infected cells. Heatmap of M_naive_ and M_IL4_ macrophage cell-surface protein expression upon stimulation with **(F)** rSFV particles alone (bottom two rows), or both virus particles and infected **(G-H)** PANC-1 or **(I-J)** Ca-Ski cancer cells. The plots represent mean value data from 4 replicates.

We further confirmed that both naïve (M_naive_) and regulatory (M_IL4_) macrophages upregulate CD80 and CD86 cell-surface expression upon stimulation with rSFV replicon particles (**Figure 7F**). Moreover, CD206 downregulation and a strong CD80 and CD86 upregulation were observed when either of the macrophage types was stimulated simultaneously with rSFV particles and infected cancer cells expressing virus-encoded IFN-γ (**Figure 7G-J**). This observation holds true for both PANC-1 (panel **G-H**) and Ca-Ski (**panel I-J**) cancer cell lines.

Additionally, we confirmed that rSFV-encoded IFN-γ mediates macrophage activation through the JAK-STAT pathway by evaluating macrophage phenotypes in the presence of the JAK inhibitor ruxolitinib. We observed that rSFV-IFN-γ, but not rSFV-Flt3L (control stimuli for myeloid cell activation) or rSFV-GFP mediated, infection of cancer cells led macrophage stimulation as measured by increased expression of co-stimulatory, anti-tumoral markers (CD80, CD86) and reduced levels of pro-tumoral markers (CD206). However, treatment with ruxolitinib attenuated these changes, indicating that IFN-γ-driven macrophage activation is at least partially dependent on JAK-STAT signaling (**Supplementary Figure 7**).

### Assessing the in vivo efficacy of rSFV-IFN-γ

Finally in a proof-of-concept experiment, we studied the effect of intratumoral administration of rSFV-IFN-γ on tumor growth and the tumor immune microenvironment (TME) in mice bearing KPC3^56–58^ pancreatic tumors (**Figure 8**). As demonstrated in (**Supplementary Figure 8**), these tumors are rich in macrophages. When tumors were palpable, mice received three intratumoral injections of PBS, rSFV-GFP, or rSFV-IFN-γ at doses of 10^7^ or 10^8^ particles on days 12, 13 and 14 post-tumor implantation (illustrated in **Figure 8A)**, Seven days after the last injection, the immune cells in tumor and spleen were analyzed by flow cytometry. Treatment with rSFV-IFN-γ at both doses markedly showed a reduction in tumor size compared to PBS and rSFV-GFP, with the highest efficacy observed at the 10^8^ dose (**Figure 8B, Supplementary Figure 9**). Notably, the higher dose of rSFV-GFP (10^8^) also led to a measurable reduction in tumor size compared to PBS, indicating that replicon treatment alone can elicit some degree of anti-tumor activity.

**Figure 8:**
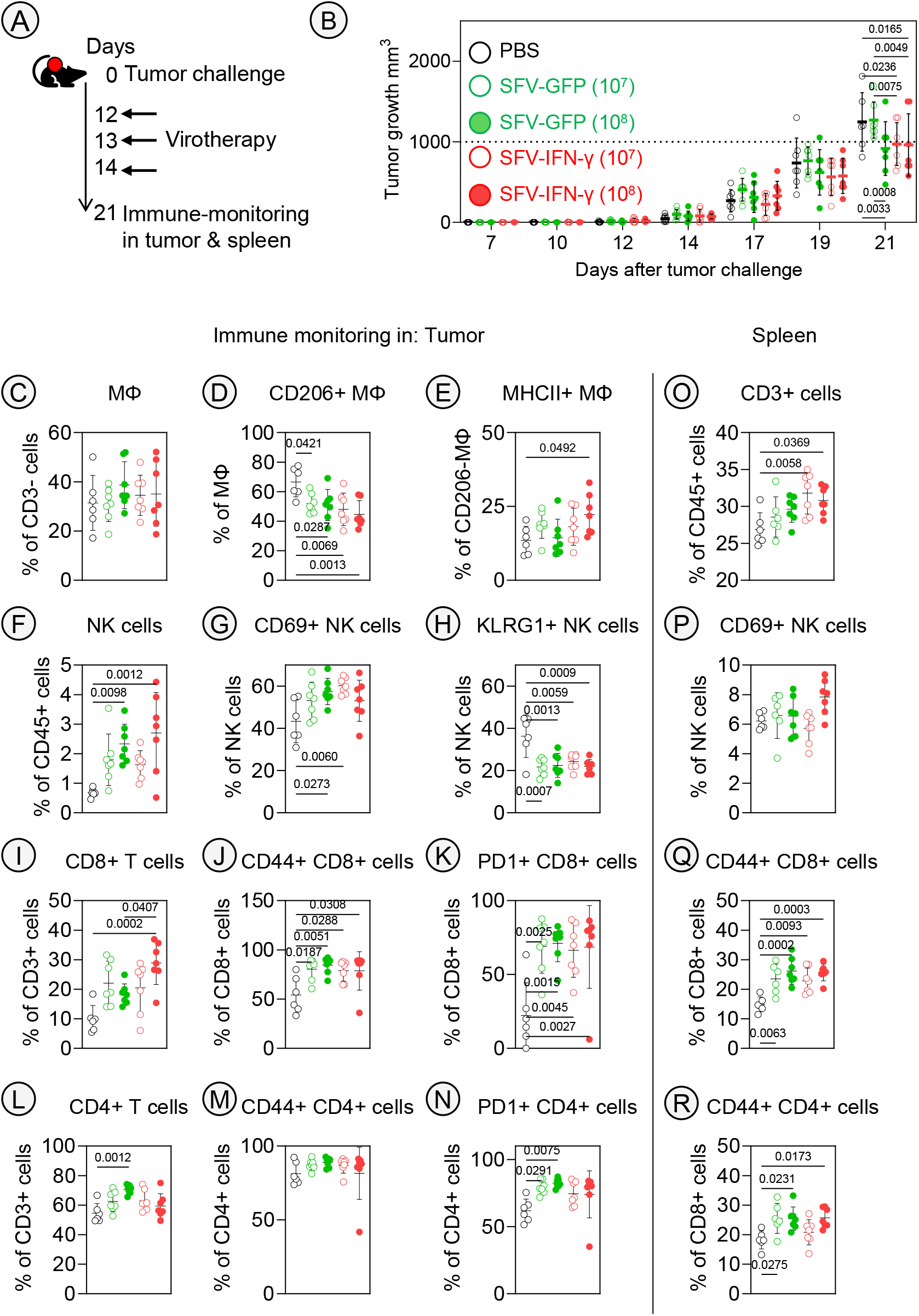
Intratumoral virotherapy with rSFV-IFN-γ induces anti-tumoral macrophage polarization and immune cell infiltration in KPC3 tumor-bearing mice. (A) C57BL/6 mice were subcutaneously injected with KPC3 tumor cells (2×10^5^) on day 0, and received intra-tumoral virotherapy with PBS (untreated control) or rSFV-replicons encoding either GFP or IFN-γ at doses of 10^7^ or 10^8^ particles per injection on days 12, 13 and 14. (B) Individual tumor growth across treatment groups, with untreated (black filled circles), SFV-GFP 10^7^ (green empty circles), SFV-GFP 10^8^ (green filled circles), SFV-IFN-γ 10^7^ (red empty circles), and SFV-IFN-γ 10^8^ (red filled circles). (C–N) Flow cytometric analysis of percentage tumor-infiltrating immune cells 7 days post-treatment: (C) total F4/80+ macrophages, (D) pro-tumoral CD206^+^ macrophages, (E) anti-tumoral MHCII+ macrophages, (F) NK cells, (G) CD69^+^ NK cells, (H) KLRG1^+^ NK cells, (I) CD8^+^ T cells, (J) CD44^+^ CD8^+^ T cells, (K) PD-1^+^ CD8^+^ T cells, (L) CD4^+^ T cells, (M) CD44^+^ CD4^+^ T cells, (N) PD-1^+^ CD4^+^ T cells. (O–R) Flow cytometric analysis of splenic immune cells: (O) CD3^+^ T cells, (P) CD69^+^ NK cells, (Q) CD44^+^ CD8^+^ T cells, (R) CD44^+^ CD4^+^ T cells. Each symbol represents an individual mouse (n=6–7 per group), with bars indicating mean ± SEM. To evaluate differences in tumor growth and immune cell infiltration following virotherapy across treatment groups, data were analyzed using one-way ANOVA to assess statistical significance, with p-value indicated per comparison. The individual tumor growth curves for respective groups are shown in **supplementary figure 9** and the flow cytometry gating strategy is illustrated in **supplementary figure 10**.

Flow cytometric analysis of tumors (**Figure 8C-N**) collected 7 days after the final injection revealed enhanced infiltration of innate and adaptive immune cells in the rSFV-IFN-γ–treated groups. Importantly, while total macrophage frequency **Figure 8C**) was not different between groups, a clear shift in macrophage polarization was observed: pro-tumoral CD206^+^ macrophages were reduced significantly (**Figure 8D, Supplementary Figure 9B**), and anti-tumoral MHCII^+^ macrophages were enriched (**Figure 8E**) in the rSFV-IFN-γ–treated tumors. These changes were dose- and stimuli dependent and most pronounced at the higher IFN-y dose. Furthermore, increased frequencies of NK cells, and CD8^+^ T-cell were found in the rSFV-IFN-γ groups compared to controls (**Figure 8F, I and L)**. NK cells and CD8+ T cells expressed more markers associated with recent antigen exposure, including CD69, CD44 and PD-1 across all rSFV-treated groups, compared to PBS-treated mice (**Figure 8 G, J, K, N**).

Second, the systemic effects were analyzed in the spleen. This analysis revealed an increased depot of CD3^+^ T-cells (Figure 8O) in mice treated specifically by the rSFV-IFN-y virus, whereas general rSFV treatment showed increased numbers of CD44+ CD4+ and CD8+ T-cell subsets (**Figure 8Q and 8R**). These data indicate that intratumoral rSFV treatment not only modified the local immune subsets in the TME, but also activated T-cells systemically.

Collectively, our data demonstrates that intratumoral administration of rSFV-IFN-γ can slow down tumor growth and modulate the TME of immune-suppressive tumors towards a more immune-infiltrated phenotype.

## Discussion

In this study, we investigated how macrophages impact the therapeutic efficacy of rSFV-based anticancer virotherapy. Our findings show that macrophages, regardless of their phenotype, do not translate rSFV-encoded proteins^37^ and simultaneously limit tumor cell infection. This highlights a significant barrier to effective virotherapy, as the presence of macrophages can limit the subsequent immunogenic activity of the virus, potentially reducing the overall therapeutic outcome^17,33^. Notably, tumor-associated macrophages often adopt immunosuppressive phenotypes that support tumor progression and contribute to an overall inhibitory tumor microenvironment. Leveraging a computational modelling approach, we hypothesized that arming therapeutic viruses to express immunogenic signals like IFN-γ could stimulate robust T-cell activation and re-polarize macrophages, tipping the balance towards tumor eradication. This was experimentally validated in both *in vitro* human-based tumor-immune cocultures and an *in vivo* murine tumor model. Overall, our results demonstrate the potential of a combined theoretical and experimental approach in improving the immunogenic potential of anticancer virotherapy.

Our computational modeling results provided insights into the spatiotemporal dynamics of rSFV-IFN-γ therapy and its potential despite an abundant presence of macrophages. Our model demonstrated that the predicted release of IFN-γ at effective concentrations is vital for achieving robust T-cell activation and subsequent tumor eradication. Specifically, the model underscored the importance of achieving the effective concentration threshold via either increasing the number of infected cells in the tumor or by improving IFN-γ production by individual infected cells; in both cases ensuring activation of anticancer cytotoxic T-cells to mount a strong antitumor response. The model also highlighted the importance of T-cell cytotoxicity in the success of rSFV-IFN-γ therapy. A minimally required cytotoxicity rate of three target cells killed per day by each cytotoxic T-cell, comparable to what is noted in the literature^38^, was found to be essential for effective tumor eradication. This emphasizes that alongside the production of IFN-γ, the intrinsic killing efficiency of T-cells is a critical factor in the overall therapeutic outcome. Importantly, the model predicted that the therapeutic success is largely independent of the number of macrophages present at the time of treatment. This suggests that rSFV-IFN-γ may effectively overcome the influence of macrophages non-permissive for SFV, provided that the thresholds for infected cells and T-cell cytotoxicity are met.

These theoretical predictions were substantiated *in vivo* using the murine KPC3 pancreatic tumor model, which is an aggressive, immunologically “cold” tumor^56–58^ known for its dense macrophage-rich barrier and poor T-cell infiltration. Intratumoral administration of rSFV-IFN-γ led to a marked delay in tumor growth compared to controls. Flow cytometry analysis of tumor-infiltrating and systemic immune cells confirmed an increased presence and activation of cytotoxic CD8^+^ T-cells, CD4^+^ T-helper cells, NK cells, and a shift in macrophage phenotype from immunosuppressive to pro-inflammatory populations. This immune reprogramming likely resulted from intra-tumoral IFN-γ expression, macrophage repolarization, or both. Notably, while rSFV-IFN-γ was most effective, even high-dose rSFV-GFP showed partial tumor growth delay, suggesting that viral infection and innate immune stimulation alone contribute to therapeutic benefit. These findings collectively underscore the potential of rSFV-IFN-γ virotherapy to enhance antitumor responses by leveraging the synergistic effects of targeted viral infection and potent immune activation, regardless of the presence of macrophages.

Based on our empirical findings, we confirm that rSFV while not infecting macrophages leads to their immunogenic activation. Importantly, as the frequency of macrophages in coculture with cancer cells increased, the infectivity of the cancer cells decreased in both monolayer and spheroid-based settings. We confirmed that macrophages themselves did not allow rSFV-encoded transgene expression, which can likely be attributed to innately active antiviral signaling pathways. This is in line with our previous work^32^ and the observations made by Olupe Kurena and colleagues^37^. Multiple oncolytic viruses have been shown to activate macrophages during therapy, both through direct virus-uptake and signaling from infected cancer cells^39–43^. Specifically for replication-competent oncolytic viruses, macrophage activation has also been attributed to therapeutic resistance due to virus-uptake by macrophages, antiviral signaling (e.g. via TNF-α or IFN pathway) and killing of infected cancer cells to inhibit virus replication^42–44^. In case of replication-deficient virotherapy like rSFV, previous studies have demonstrated that sensing of viral RNA by various endosomal toll-like receptors (TLRs) and cytosolic retinoic acid-inducible gene I (RIG-I) receptor can result in degradation of viral RNA to downregulate viral-protein expression in macrophages^9^. However, TLR and RIG-I mediated sensing also may lead to the proinflammatory activation of macrophages^45,46^. In line with our previous study^32^, we observed that macrophages upregulate CD80 and CD86 surface protein expression upon viral stimulation, corresponding to a stronger co-stimulatory signaling for antigen presentation. This is attributed partially via activation of JAK-STAT and NF-κB signaling in macrophages^32^. A similar macrophage activation profile was observed independent of their initial naïve (M_naive_) or regulatory (M_IL4_) macrophage phenotype. Various groups have demonstrated so far that although macrophages demonstrate functionally distinct phenotypes^47,48^, they are capable of switching through these phenotypes in response to environmental stimuli within 48 hours^49,50^. Despite their adaptability, macrophages maintain their non-permissive nature towards rSFV-mediated transgene expression independent of their initial phenotype, their expression of rSFV-entry receptors, and their switch towards a pro-inflammatory phenotype upon rSFV-mediated stimulation.

Engineering rSFV to encode IFN-γ in infected cancer cells enhanced macrophage polarization to a proinflammatory phenotype, is consistent with the increased frequency of anti-tumoral macrophages observed *in vivo* following rSFV-IFN-γ treatment. This aligns with previous findings where rSFV driven IFN-γ expression induced a strong antitumor immune response and inhibited tumor growth in a murine orthotopic breast cancer model, by promoting macrophage activation and remodeling the tumor microenvironment^51^. In our study, macrophages upregulated pro-inflammatory markers (CD80, CD86) and downregulated pro-tumoral markers (CD206) to a significantly higher magnitude when in the presence of infected cancer cells expressing rSFV-encoded IFN-γ, as compared to being stimulated by virus alone or rSFV-GFP and rSFV-Flt3L infected cancer cells, where Flt3L served as an myeloid-stimulating control. We assume this to be a result of the bimodal stimulation by virus-particles and extracellular sensing of IFN-γ. Previous studies have demonstrated that either type-I or type-II IFNs are required for macrophages to polarize towards a strongly pro-inflammatory phenotype when stimulated by stress signals like extracellular release of heat-shock proteins or danger signal like bacterial lipopolysaccharides and viral RNA^48,49,52^. Therefore, it is likely that the intracellular release of rSFV-RNA upon virus uptake in the macrophages and extracellular IFN-γ produced by neighboring infected cancer cells results in their proinflammatory activation. We indeed confirmed that macrophage activation by rSFV-encoded IFN-γ can be reversed upon blocking JAK-STAT signaling. Furthermore, as we did not observe rSFV-mediated transgene expression by macrophages, we consider that rSFV-encoded IFN-γ expression is limited to infected cancer cells and that IFN-γ is not sensed in an autocrine manner by macrophages.

Our study demonstrates that rSFV-encoding IFN-γ significantly stimulates T-cell responses, independent of the influence of macrophages. This was observed across both monolayer (2D) and spheroid (3D) cocultures, as well as *in vivo* highlighting the robustness of this approach. In monolayer cocultures, the presence of rSFV-IFN-γ led to pronounced activation of both CD4 and CD8 T-cells, even in the presence of high macrophage frequencies. Here, T-cell activation relied on IFN-γ or macrophage polarization and was diminished upon inhibition of JAK-STAT signaling. Notably, macrophages, despite their non-permissiveness to SFV infection and potential for antiviral signaling, did not impede T-cell activation induced by rSFV-IFN-γ. This finding underscores the potential of IFN-γ as a powerful immunostimulatory cytokine that can override the suppressive effects of macrophages in the tumor microenvironment. Furthermore, the impact of macrophage phenotype and frequency on T-cell responses varied between monolayer and spheroid cocultures. In monolayer systems, macrophages reduced CD4 but not CD8 T-cell activation, suggesting a differential regulatory effect on helper versus cytotoxic T-cells. However, in spheroid cocultures, an increase in macrophage frequency led to a reduction in activation of both CD4 and CD8 T-cells. Despite these variations, rSFV-IFN-γ consistently promoted strong T-cell responses regardless of either a naïve (M_naive_) or regulatory (M_IL4_) macrophage phenotype. Finally, in both monolayer and spheroid cocultures, a high frequency of macrophages correlated to a stronger CD4 T-cell activation by rSFV-IFN-γ, which may be attributed to rSFV-mediated proinflammatory activation of the macrophages. In addition to transgenic IFN-γ expression, differences in innate cancer cell-mediated immune signaling were also found to influence both macrophage and T-cell activation, as noted in a previous study^23^. Nevertheless, IFN-γ expression in infected PANC-1 and Ca-Ski cells proved effective in inducing robust immune responses. Observations of Meissner et al. corroborate our results, that encoding IFN-γ by a non-replicating virotherapy, influenza A virus in their case, can promote tumor eradication through enhanced immune responses and not primarily through virus-mediated oncolysis^53^. Indeed, a replication-competent virotherapy causing extensive virus-mediated oncolysis in combination with IFN-γ mediated immune stimulation can also lead to tumor eradication albeit with a risk of virus persistence^54^.

In conclusion, our study presents a compelling case for the use of rSFV-encoding IFN-γ as a means to enhance antitumor immune responses, effectively overcoming macrophage-mediated regulation of virus-infection and T-cell activation. The combination of computational modeling with experimental validation in diverse human tumor models, provides evidence supporting this therapeutic strategy. Future studies should explore the clinical translation of these findings, assessing the efficacy of rSFV-IFN-γ in preclinical models like patient-derived organoids to test its therapeutic potential in a patient-specific manner. The significant implications of our findings lie in their potential to inform the design of anticancer virotherapies, especially for tumors characterized by high macrophage infiltration and immune suppression.

### Limitations of the study

The primary aim of our study was to assess and improve the efficacy of rSFV replicons in the context of tumor-associated macrophages in a human model. While our study demonstrates the potential of rSFV-encoding IFN-γ in enhancing T-cell immune responses, several limitations must be acknowledged. One significant limitation of our study is the lack of direct measurement of T-cell-mediated tumor cell killing *in vitro*, due to the unavailability of a tumor-specific T-cell model. However, the observed tumor growth inhibition and increased immune cell infiltration *in vivo* following rSFV-IFN-γ treatment provide indirect evidence supporting the functional relevance of the T-cell activation seen in our coculture systems. Furthermore, our study does not fully account for the complex interactions within the tumor immune microenvironment, focusing primarily on T-cell responses. This narrow focus may overlook the broader spectrum of immune cell dynamics and the potential contributions of other immune cells, such as dendritic cells. Additionally, the *in vitro* coculture models, while informative, do not fully replicate the *in vivo* human tumor microenvironment. Expanding the study to test different virotherapies in the context of *in vivo* humanized murine models with xenograft tumors or patient-derived organoids representing diverse tumor and HLA-types will be essential to validate these findings and to fully understand the therapeutic potential of immunogenic virotherapy in a clinical setting.

## Acknowledgements

This study was partially funded by Stichting De Cock-Hadders (grant 2020-35) awarded to DB and by Stichting Overleven met alvleesklierkanker (grant SOAK 22.02) awarded to NvM. Part of the work has been performed at the UMCG Imaging and Microscopy Center (UMIC) and the Flow Cytometry Research Unit (FCU) which is sponsored by UMCG. We thank the Center for Information Technology of the University of Groningen for their support and for providing access to the Hábrók high-performance computing cluster. We acknowledge the Flow Cytometry Facility and animal facility at LUMC for their support with the *in vivo* study and immunological analyses.

## Author contributions

All authors made substantial contributions to the manuscript.

**LH:** Writing and editing manuscript, Data collection and analysis, Validation, Visualization

**PK:** Writing and editing manuscript, Data collection and analysis, Validation, Visualization

**TJ:** Conceptualization, Writing and editing manuscript, Software, Data analysis, Visualization, Supervision

**BNH:** Writing and editing manuscript, Data collection and analysis, Validation, Visualization

**FJW:** Conceptualization, Writing and editing manuscript, Supervision

**NM**: Conceptualization, Writing and editing manuscript, Supervision, Funding acquisition

**TD:** Conceptualization, Writing and editing manuscript, Supervision, Funding acquisition

**DB:** Conceptualization, Writing and editing manuscript, Data collection and analysis, Software, Visualization, Supervision, Funding acquisition

## STAR Methods

### Key resources table

| REAGENT or RESOURCE | SOURCE | IDENTIFIER |
| --- | --- | --- |
| Antibodies |  |  |
| anti-human CD4 (OKT4) FITC | Biolegend | 317408 |
| anti-human CD8 (SK11) APC-Cy7 | Biolegend | 344714 |
| anti-human CD38 (HIT2) BV421 | Biolegend | 303526 |
| anti-human CD69 (FN50) APC | Biolegend | 310910 |
| anti-human CD107a (H4A3) PE | Biolegend | 328608 |
| anti-human CD223 (LAG-3) (11C3C65) PE | Biolegend | 369306 |
| anti-human CD152 (CTLA-4) (BNI3) BV421 | Biolegend | 369606 |
| anti-human CD279 (PD-1) (A17188B) APC | Biolegend | 621610 |
| anti-human CD11b (ICRF44) PE | Biolegend | 301306 |
| anti-human CD206 (15-2) APC-Fire | Biolegend | 321133 |
| anti-human CD80 (2D10) APC | Biolegend | 374204 |
| anti-human CD86 (BU63) FITC | Biolegend | 328608 |
| anti-human CD163 (GHI/61) PE | Biolegend | 333605 |
| anti-human CD68 (Y1/82A) PE-Cy7 | Biolegend | 333815 |
| anti-human CD206 (MMR) APC | Biolegend | 321109 |
| anti-human CD80 (2D10) BV421 | Biolegend | 305221 |
| anti-murine CD45.2 (104) FITC | Biolegend | 109806 |
| anti murine CD45.2 (104) AF700 | Biolegend | 109822 |
| anti-murine CD3 (145-2C11) PE-CF594 | BD Biosciences | 562286 |
| anti-murine CD8α (53-6.7) AF700 | eBioscience | 56-0081-82 |
| anti-murine NK1.1 (PK136) BV650 | BD Biosciences | 564143 |
| anti-murine CD4 (RM4-5) APC | BioLegend | 100547 |
| anti-murine F4/80 (BM8) PE-Cy7 | Biolegend | 123114 |
| anti-murine CD206 (C068C2) BV711 | Biolegend | 141727 |
| anti-murine Ly6G (1A8) BV786 | Biolegend | 127645 |
| anti murine Ly6C (HK1.4) BV605 | Biolegend | 128036 |
| Anti mouse/human CD44 (IM7) BV785 | Biolegend | 103059 |
| Anti mouse CD69 (H1.2F4) FITC | eBioscience | 11-0691-82 |
| Anti mouse KLRG1 PE-Cy7 | eBioscience | 25-5893-82 |
| Anti mouse CD279 (PD-1) (28F.1A12) PE | Biolegend | 135205 |
| anti-murine MHC-II (M5/114) BV421 | BD Biosciences | 562564 |
| Zombie aqua viability dye | Biolegend | 423102 |
| Bacterial and virus strains |  |  |
| Plasmid (Helper-2 and SFV-transgene) constructs to produce recombinant SFV replicon particles | Liljestrom lab, Karolinska Institutet Sweden | NA |
| E coli K12 JM110 | Agilent | 200239 |
| rSFV-GFP | This study (in-house) | NA |
| rSFV-IFN-γ | This study (in-house) | NA |
| Biological samples |  |  |
| Human buffy coats for PBMC isolation from HLA-A*02:01-typed healthy donors | Sanquin, Nijmegen, The Netherlands | NA |
| Chemicals, peptides, and recombinant proteins |  |  |
| Restriction enzyme PspOMI | ThermoFischer Scientific | ER0131 |
| Restriction enzyme XmaI | ThermoFischer Scientific | ER0171 |
| Restriction enzyme SpeI | ThermoFischer Scientific | ER1251 |
| SP6 polymerase | Amersham Pharmacia Biotech, Piscataway, US | NA |
| Celltrace FarRed dye | Invitrogen | C34572 |
| Ruxolitinib | Stem Cell Technologies | 73402 |
| α-chymotrypsin | Sigma Aldrich | C4129 |
| Aprotinin | Sigma Aldrich | A1153 |
| Macrophage colony stimulating factor (M-CSF) | Biolegend | 572903 |
| Interleukin-4 (IL-4) | Biolegend | 574004 |
| Interferon-gamma (IFN-γ) | Biolegend | 570204 |
| Lipopolysaccharides (LPS) from *E. coli* O128:B12 | Sigma Aldrich | L2755 |
| LIberase TL | Sigma Aldrich | 05401020001 |
| Mouse BD Fc block (anti-mouse CD16 and CD32) | BD Biosciences | 553142 |
| Hydrogen peroxidase | VWR | 7422478 |
| ELK powder | Campina | NA |
| Rat, anti-mouse F4/80 antibody | Sanbio | CI:A3-1 |
| Biotinylated rabbit anti-rat antibody | Agilent | NA |
| VECTASTAIN Elite ABC HRP Kit | Vector laboratories | PK-6100 |
| Entellan | Sigma Aldrich | 107960 |
| Critical commercial assays |  |  |
| NA |  |  |
| Deposited data |  |  |
| NA |  |  |
| Experimental models: Cell lines |  |  |
| Hamster, BHK-21 cells | ATCC | ccl-10 |
| Human, PANC-1 cells | ATCC | crl-1469 |
| Human, Ca-Ski | ATCC | crl-1550 |
| Murine, KPC3 cells | Nadine van Montfoort lab, Leiden University Medical Center, Leiden | RRID:CVCL_A9ZK |
| Experimental models: Organisms/strains |  |  |
| C57BL/6J mice | Charles River Laboratories | C57BL/6NCrl |
| Oligonucleotides |  |  |
| NA |  |  |
| Recombinant DNA |  |  |
| Human IFN-gamma gene | This paper | NA |
| Murine IFN-gamma gene | This paper | NA |
| Software and algorithms |  |  |
| Graphpad Prism | Graphpad | https://www.graphpad.com/ |
| Incucyte® Software Modules | Sartorius | https://www.sartorius.com/ |
| Qpath | Qpath | https://qupath.github.io/ |
| FlowJo | FlowJo | https://www.flowjo.com/ |
| Rstudio | Posit (Rstudio) | http://www.posit.co/ |
| RawGraphs | RawGraphs | https://www.rawgraphs.io/ |
| Other |  |  |
| Nunclon Sphera 96 well plates | ThermoFischer Scientific | 174925 |
| Oncolytic Virus Immune simulator (OVI) model | This paper | https://github.com/rugtres/OVI/tree/macrophage |
| Superfrost Plus slides | VWR | 48311-703 |

## Resource availability

### Lead contact

Further information and requests for resources and reagents should be directed to and will be fulfilled by the lead contact Darshak K. Bhatt

### Material availability

All unique/stable reagents generated in the study are available upon request, except where restricted by institutional agreements or MTAs.

### Data and code availability

- All data reported in this paper will be shared by the lead contact upon request.
- Any additional information required to reanalyze the data reported in this paper is available from the lead contact upon request.
- The code used for this work and an executable version of the Oncolytic Virus Immune simulator (OVI) can be found at https://github.com/rugtres/OVI/tree/macrophage.

## Method details

### TCGA analysis

We used the online CRI iAtlas portal to analyze TCGA data of cervical and pancreatic cancer patients^1,36^. We selected the TCGA cohort of cervical cancer (CESC) patients (n=300) and pancreatic cancer (PAAD) patients (n=151). Performed Pearson correlation to analyze the relationship between IFN-γ signature, cytotoxic CD8 T-cells, and pro-tumoral (M2-like or M_alter_) or anti-tumoral (M1-like or M_class_) macrophages. IFN-γ signature and immune cell subset quantifications of the tumor samples were performed previously as a part of the immune landscape analysis of cancer^1,36^.

### Computational model

In our previous work^33,34^, we developed a spatiotemporal model for the effect of anticancer virotherapy on tumor development to evaluate how anticancer T-cell responses triggered by immunogenic signals affect the therapeutic outcome. In the current study, we employed the model to simulate a tumor environment with three cell types: stromal macrophages, uninfected infection-sensitive cancer cells, and infected cancer cells. We assumed that infection-sensitive cancer cells and stromal macrophages are initially distributed randomly. Cells can divide, change status, or die, with varying rates across cell types. Cancer cells can be infected by a therapeutic virus, which targets and kills them, while stromal macrophages are non-permissive to infection. Since rSFV is a non-replicating virotherapy, no further viral spread occurs in the model. We incorporated a cell-specific immune response to assess how virus-induced immunogenic signals influence therapeutic outcomes. The model assumes that infection-induced cell death induces the release of immunogenic signals, which diffuses and subsequently activates T-cells to kill cancer cells, with cytotoxicity of the activated T-cells depending on the local concentration of immunogenic signals. Key variables assessed in this study include the percentage of infected cancer cells, amount of immunogenic signal released per infected cell, T-cell cytotoxicity rate, and macrophage frequency. Immunogenic molecules are initially absent and increase in concentration (λ) in the grid cell upon infection-induced cell death. Immunogenic signals disperse to neighboring cells via diffusion and diminishes over time due to evaporation. In our previous studies, the model revealed that therapeutic outcomes are probabilistic. To account for this, we ran 10,000 simulations per parameter combination to capture stochastic variations in the results. Since rSFV is a suicidal virotherapy and does not replicate, the model predicted two possible outcomes at 1,000 days: (i) total tumor eradication, where all cancer cells are eliminated; or (ii) tumor persistence, where cancer cells survive due to insufficient virus- and/or immune-mediated killing. For this study, we focused on the probability of total tumor eradication as the primary outcome. A detailed description of the model can be found in references 33 and 34. The code used for this work and an executable version of the Oncolytic Virus Immune simulator (OVI) can be found at https://github.com/rugtres/OVI/tree/macrophage.

### Cell culture

BHK-21, Ca-Ski, and Panc-1 cells were cultured in RPMI 1640 medium supplemented with 10% fetal bovine serum (FBS), 100 U/ml penicillin, and 100 µg/ml streptomycin (P/S, Thermo Scientific). Since, PANC-1 and Ca-Ski cells are HLA-A*02:01-restricted, we performed the cancer-immune coculture assays with peripheral blood mononuclear cells (PBMCs) derived from HLA-A*02:01-matched healthy donors (Sanquin, Netherlands). Cancer cell-macrophage cocultures were cultured in RPMI 1640 medium supplemented with 10% FBS and 100 U/ml penicillin, 100 µg/ml streptomycin. Cocultures with PBMCs were similarly cultured in RPMI 1640 medium supplemented with 10% FBS and 100 U/ml penicillin, 100 µg/ml streptomycin. For spheroid cocultures, cells were seeded in NuncSphera round-bottom 96-well plates (Thermo Fischer) and centrifuged for 10 minutes at 1500 RPM. All cells were maintained in an incubator at 37°C with 5% CO_2_.

### Monocyte derived macrophage differentiation and polarization

Peripheral blood derived monocytes were differentiated to macrophages and polarized in RPMI 1640 medium supplemented with 10% FBS, 100 U/ml penicillin, 100 µg/ml streptomycin, 1mM sodium pyruvate (Thermo Scientific), non-essential amino-acids (NEAA, Thermo Fisher), and 100 ng/mL macrophage colony-stimulating factor (M-CSF, BioLegend San Diego, CA, USA). Briefly, PBMCs from healthy donors were cultured for 2 hours to allow monocyte adherence. Afterward, non-adherent cells were gently removed. The adhered cells were stimulated with medium containing 100 ng/ml M-CSF. The medium containing M-CSF was refreshed on 3 days post culture. Macrophages were replated on 7 days of culture. 24 hours later, macrophages were polarized to either a M_IL4_ state with medium containing 20 ng/mL IL-4 (BioLegend, San Diego, CA, USA), or a M1-like state with medium containing 20 ng/ml IFN-γ and 100 ng/ml lipopolysaccharides (LPS), or were kept unstimulated (M_naive_).

### Design, production and titer determination of rSFV

We have previously designed rSFV-replicon particles expressing immunogenic transgenes and capable of a single round of infection^23,55^. Briefly, enhanced green fluorescent protein (GFP) and IFN-γ (human and murine versions) as transgenes were ordered as a DNA construct (Eurofins Genomics, Ebensburg, Germany) and were cloned in the SFV-replicon backbone plasmid (pSFV) by using PspOMI and XmaI as restriction sites and *E. coli* JM110 as the competent cell chassis. Sanger sequencing was performed on isolated clones to validate insertion (Eurofins Genomics). pSFV containing either GFP or IFN-γ and a SFV-Helper-2 plasmid (hSFV) were linearized with SpeI digestion (Life Technologies) for *in vitro* RNA synthesis by SP6 polymerase reaction (Amersham Pharmacia Biotech, Piscataway, US). Next, pSFV-transgene RNA and hSFV RNA were mixed in a 2:1 ratio and co-transfected in BHK21 cells in the presence of electroporation buffer using the BioRad Gene Pulser II system (2 pulses, 850 V/25 µF; Biorad, Hercules, U.S.A.). After electroporation, the cells were cultured in RPMI 1640 media supplemented with 5% FBS, 100 U/ml penicillin, and 100 µg/ml streptomycin for 48 hours at 30°C with 5% CO_2_. The rSFV-transgene particles were purified by discontinuous sucrose density gradient ultracentrifugation and stored in TNE buffer as aliquots at -80°C. Before use, all rSFV particles were activated by the addition of 1:20 volume 10 mg/ml α-chymotrypsin (Sigma Chemical, St. Louis, US) and 2 mM CaCl_2_ for 30 minutes to cleave the mutated spike proteins. After which, the α-chymotrypsin was inactivated by the addition of 1:2 volume 2 mg/ml aprotinin (Sigma Chemical). Finally, the titer determination of rSFV-particles was performed as described previously. Briefly, rSFV-particles were titrated by serial dilution on monolayers of BHK-21 cells cultured in LabTek slides. After infection and incubation for 24 hours, the cells were fixed in 10% (w/v) acetone and further stained for nsP3 using a primary polyclonal rabbit-anti-nsP3 antibody (1:2000 dilution), whilst a secondary Cy3-labeled animal-anti-rabbit antibody (1:200 dilution) was used to amplify the signal. Positive cells were counted using fluorescence microscopy, and the titers were determined.

### Evaluating virus-infection in macrophage-cancer cell cocultures

M_naive_ or M_IL4_ macrophages were differentiated from human PBMCs as described above. Different frequencies of M0 or M_IL4_ macrophages were cocultured with either a constant or variable frequency of Panc-1 or Ca-Ski cancer cells. When specified, macrophages were stained with CellTrace™ Far Red dye (ThermoFischer Scientific, C34564) following the recommended protocol from the manufacturer. After overnight incubation, cocultures were infected with an MOI of 10 (multiplicity-of-infection) of rSFV-GFP. The ability of rSFV-particles to infect cells and express GFP was monitored over 24 hours by Incucyte-based brightfield and fluorescence microscopy. The number of GFP-positive cells served as a measure of the rSFV-GFP particle’s infectivity.

### Evaluating macrophage activation by virus particles and infected cancer cells

M_naive_ or M_IL4_ macrophages were differentiated from human PBMCs as described above. M_naive_ or M_IL4_ macrophages were cocultured in a 1:2 ratio with Panc-1 or Ca-Ski cells in a 24-well plate. After overnight incubation, cocultures were infected with MOI-10 of rSFV-GFP or rSFV-IFN-γ. 24 hours post-infection (HPI), all cells were collected and processed for Flow cytometry-based analysis of macrophage activation markers. The gating strategy is illustrated in **Supplementary Figure 7**.

### Evaluating T-cell activation in tumor-immune cocultures

M_naive_ or M_IL4_ macrophages were differentiated as described above, from HLA-A*02:01 typed healthy donors. They were cocultured with Panc-1 cells or Ca-Ski cells for 24 hours in 2 tumor cells to 1 macrophage cell ratio (15 000 cells per well) in a treated 96-well plate. Cocultures were then infected with MOI-10 of rSFV-particles encoding GFP or IFN-γ. 8 hours post-infection, freshly thawed PBMCs (from the same donor as used for generating macrophages) were added to the cocultures (75 000 cells per well). 18 hours post PBMCs addition, all cells were collected and stained for CD4 and CD8 T-cell population, as well as immune activation cell surface markers, and analyzed *via* Flow cytometry. The gating strategy is illustrated in **Supplementary Figure 4**.

### Animal experiments

*In vivo* experiments were conducted as described previously^56–58^. In brief, male C57BL/6J mice (6–8 weeks old; were obtained from Charles River Laboratories and housed in individually ventilated cages (3-4 mice/cage) at the LUMC animal facility. The experiment was reviewed, ethically approved, and registered by the institutional Animal Welfare Body of Leiden University Medical Center. The experiment was carried out in compliance with Dutch and EU regulations on animal experimentation under project license AVD1160020187004, issued by the competent authority on animal experiments in the Netherlands. After one week of acclimatization, mice were subcutaneously injected in the right flank with 2×10^5^ KPC3 cells in 100 μL PBS/0.1% BSA. Once tumors became palpable, mice were stratified into groups with similar average tumor volumes and treated intratumorally under isoflurane anesthesia with 10^7^ or 10^8^ infectious particles of rSFV-GFP, rSFV-IFN-γ, or PBS (30 μL) on three consecutive days. In contrast to the previous experiments, rSFV-encoding murine-IFN-γ was used. Tumor volume, welfare and body weight were monitored thrice weekly using calipers (tumor volume = length × width × height), blinded when possible. Mice were sacrificed for immune profiling 7 days post-treatment or earlier if they reached human endpoint, as described to be tumors exceeding 1000mm^3^, signs of tumor ulceration or non-experimental related complications (n=1 mouse, that was excluded due to fighting wounds). Tumors were digested with Liberase TL for 15 min at 37°C, and processed into single-cell suspensions. Cells were stained with Zombie Aqua viability dye, incubated with Mouse BD Fc Block™ (anti-Mouse CD16/CD32), stained for cell surface markers and fixed in 1% paraformaldehyde. Flow cytometry was performed on a BD LSRFortessa X20 at the LUMC Flow Cytometry Facility and analyzed using FlowJo v10. The gating strategy is illustrated in **Supplementary Figure 10**.

### Immunohistochemistry staining

Formalin fixed tumors were embedded in paraffin and sectioned at 4μm before placing on Superfrost^®^ Plus slides (VWR). Sections were dried overnight at 37°C and stored at 4°C until staining. Slides were deparaffinized and endogenous peroxidase was blocked with 0,3% hydrogen peroxidase (VWR) in methanol for 20 minutes. Rehydration was followed by 16 minute boiling in Trypsin/Calcium chloride solution (pH 7,4) followed by blocking with 0,2% ELK (Campina) in PBS/0,1% BSA. Overnight incubation with primary rat anti-mouse F4/80 (CI:A3-1, Sanbio) was performed at RT in humidified box. The next morning, samples were incubated for 30 minutes at RT with biotinylated rabbit anti-rat (Agilent), followed by 30 minutes incubation with avidin-biotin complex (VECTASTAIN^®^ Elite^®^ ABC HRP Kit; Vector Laboratories). Peroxidase activity was detected using the 2-component liquid DAB+ system (Agilent) according to the manufacturer’s instructions for 5 min. Counterstaining was performed using hematoxylin (Sigma Aldrich) for 15 seconds, followed by dehydration and mounting using Entellan (Sigma Aldrich). Slides were scanned using Zeiss Axio Scan Z.1 and processed with QuPath.

### Quantification and statistical analysis

Experimental data represents the mean ± SEM of the number of replicates. Statistical analysis of the data is specified in the respective figure legends when applicable. For linear correlation, bivariate linear regression was performed, with r (correlation coefficient) and p-values indicated per plot. one-way ANOVA was also used to compare groups and assess statistical significance, with p-value indicated in the plots per comparison. Graphs were made using Rstudio, RawGraphs, and Graphpad Prism 10.

## Supplementary file

All the supplementary figures can be found in the supplementary document.

