## Supplementary figures for "Alphavirus replicons encoding IFN-γ enhance cancer virotherapy by overcoming macrophage-mediated suppression"

### Supplementary Figure 1

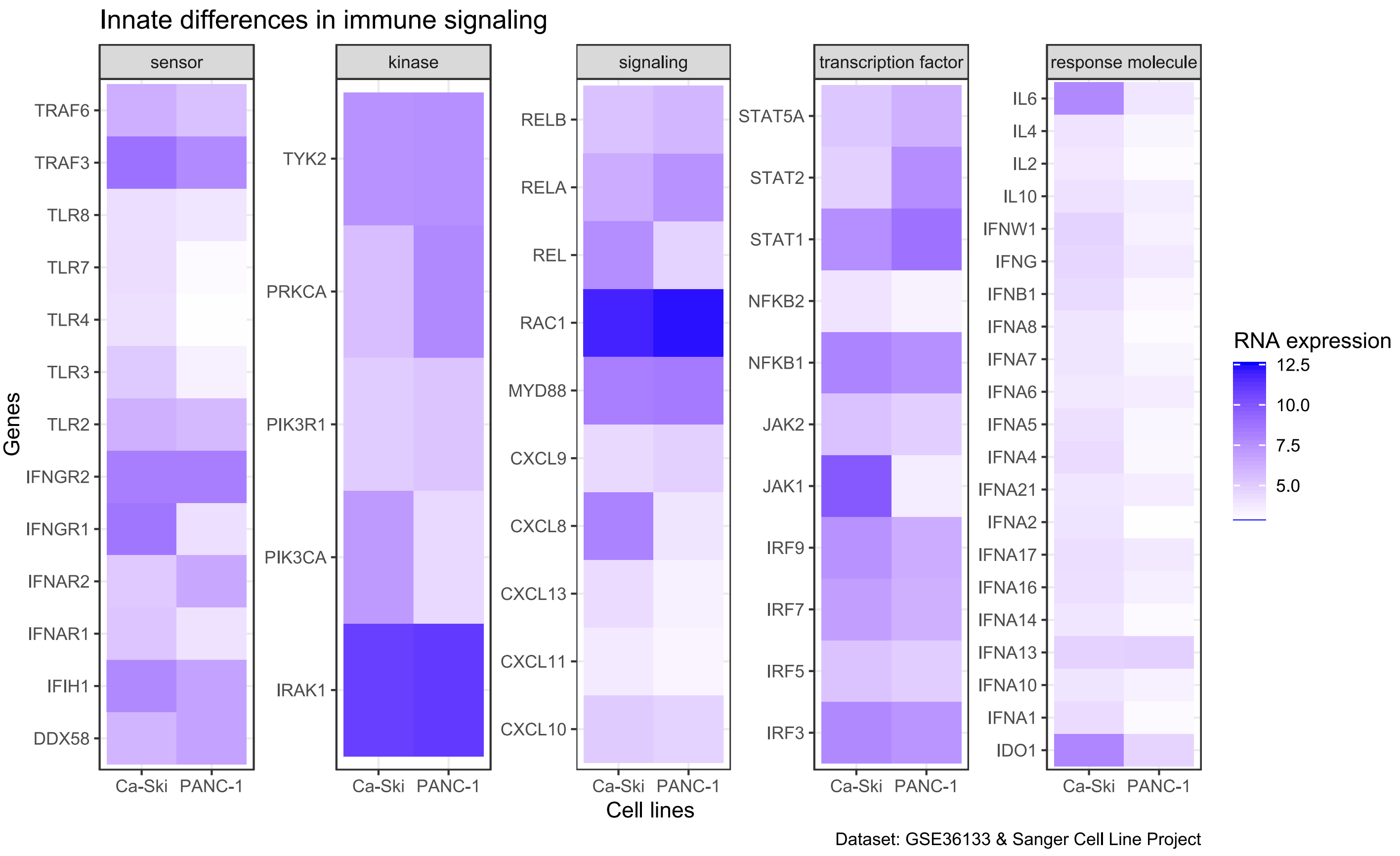


**Supplementary Figure 1:** Innate differences in RNA expression of genes involved in immune signaling pathways between Ca-Ski and PANC-1 cell lines. Gene expression data (calculated RNA expression) were obtained from the GSE36133 and Sanger Cell Line Project datasets. Genes are categorized into five functional groups: *sensor, kinase, signaling, transcription factor,* and *response molecule*. Differences in expression levels between the two cell lines highlight potential variations in innate immune signaling activity. Darker blue indicates higher RNA expression.

**Supplementary Figure 2**

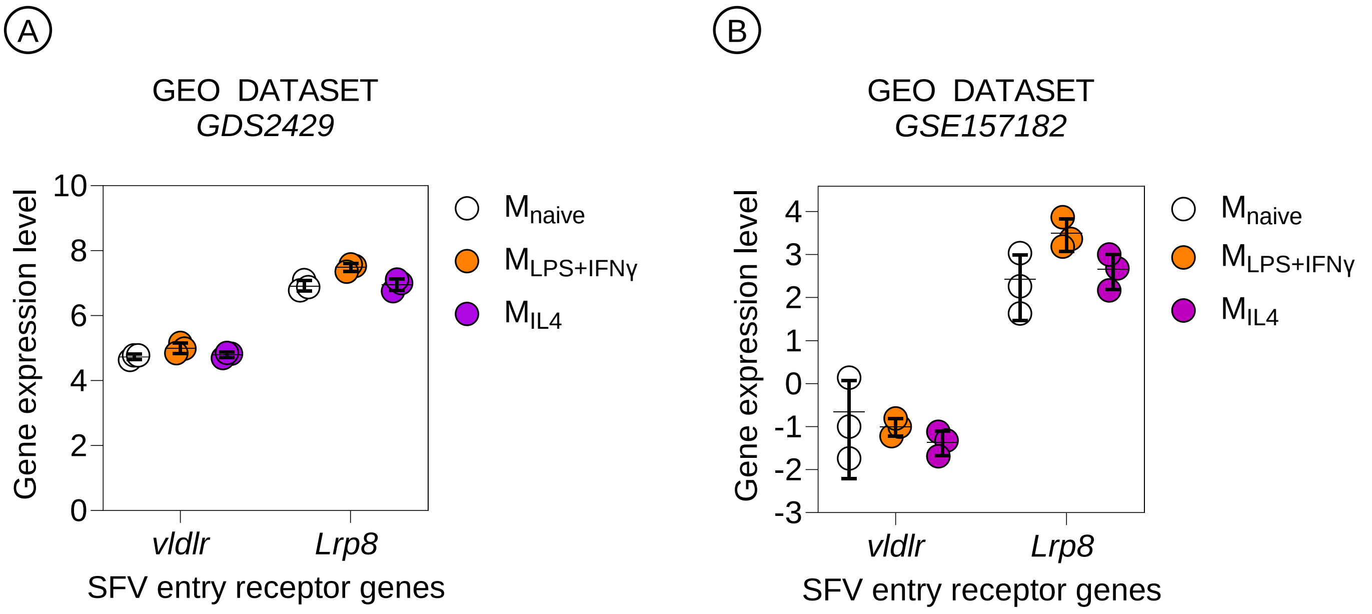


**Supplementary Figure 2:** RNA expression of genes coding for SFV-entry receptors in peripheral blood monocyte-derived macrophages polarized to different phenotypes. The data is analyzed from the GEO DATASETs (A) GDS2429 and (B) GSE157182, and is represented as mean values with standard deviation with 3 replicates per condition.

### Supplementary Figure 3

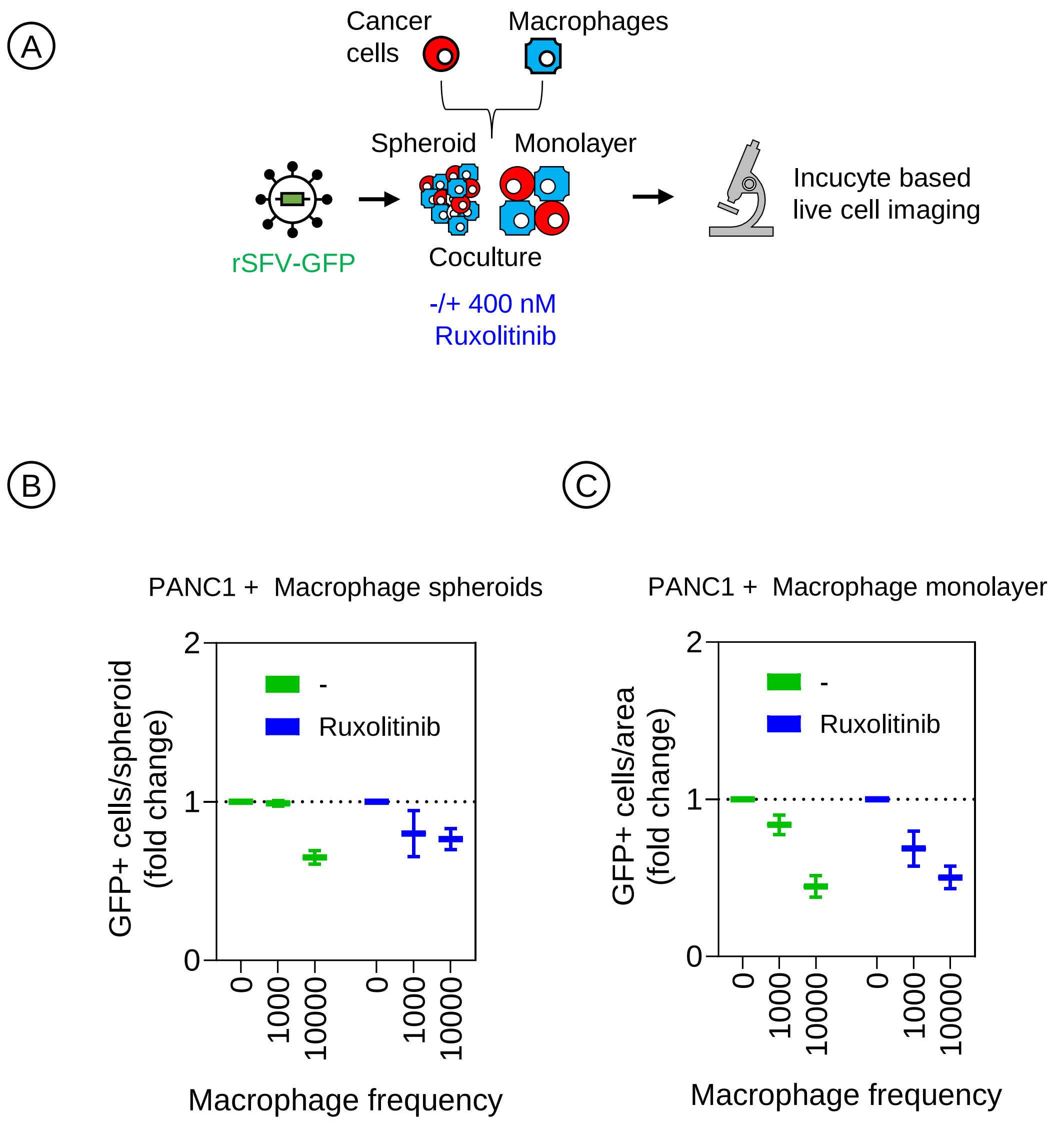


**Supplementary Figure 3:** Effect of JAK-STAT pathway inhibition on cancer cell infection in presence of macrophages. (A) The experimental setup is the same as described in Figure 1, with the addition of treatment with a JAK-STAT inhibitor (400nM Ruxolitinib). Quantitative analysis of GFP expression indicating virus infection in the (B) spheroid and (C) monolayer coculture of PANC-1 cells with $M_{IL4}$ macrophages in the presence or absence of Ruxolitinib. The plots represent data from 4 replicates of respective coculture methods. Data are presented as fold change mean values ± SD.

### Supplementary Figure 4

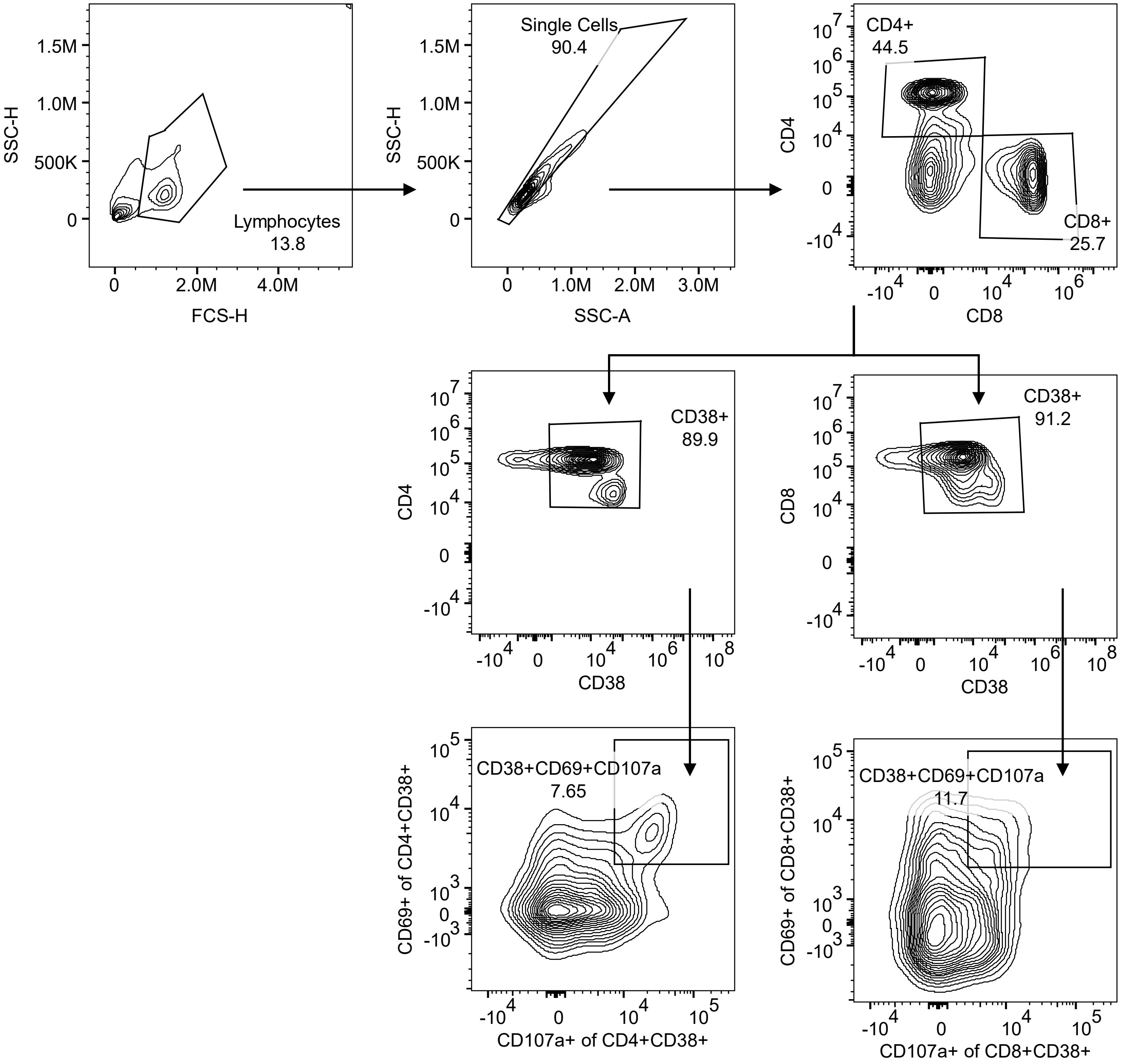


**Supplementary Figure 4:** Flow cytometry gating strategy for identifying activated CD4+ and CD8+ T cells expressing CD38, CD69, and CD107a. (Top row) Lymphocytes were first gated based on forward and side scatter properties, followed by gating on single cells. T cell subsets were identified based on CD4 and CD8 expression. (Middle row) CD4+ and CD8+ T cells were further gated for CD38 expression. (Bottom row) CD38+ T cells were analyzed for co-expression of CD69 and CD107a, identifying highly activated or cytotoxic T cell populations. Percentages represent the proportion of cells within each gate.

**Supplementary Figure 5**

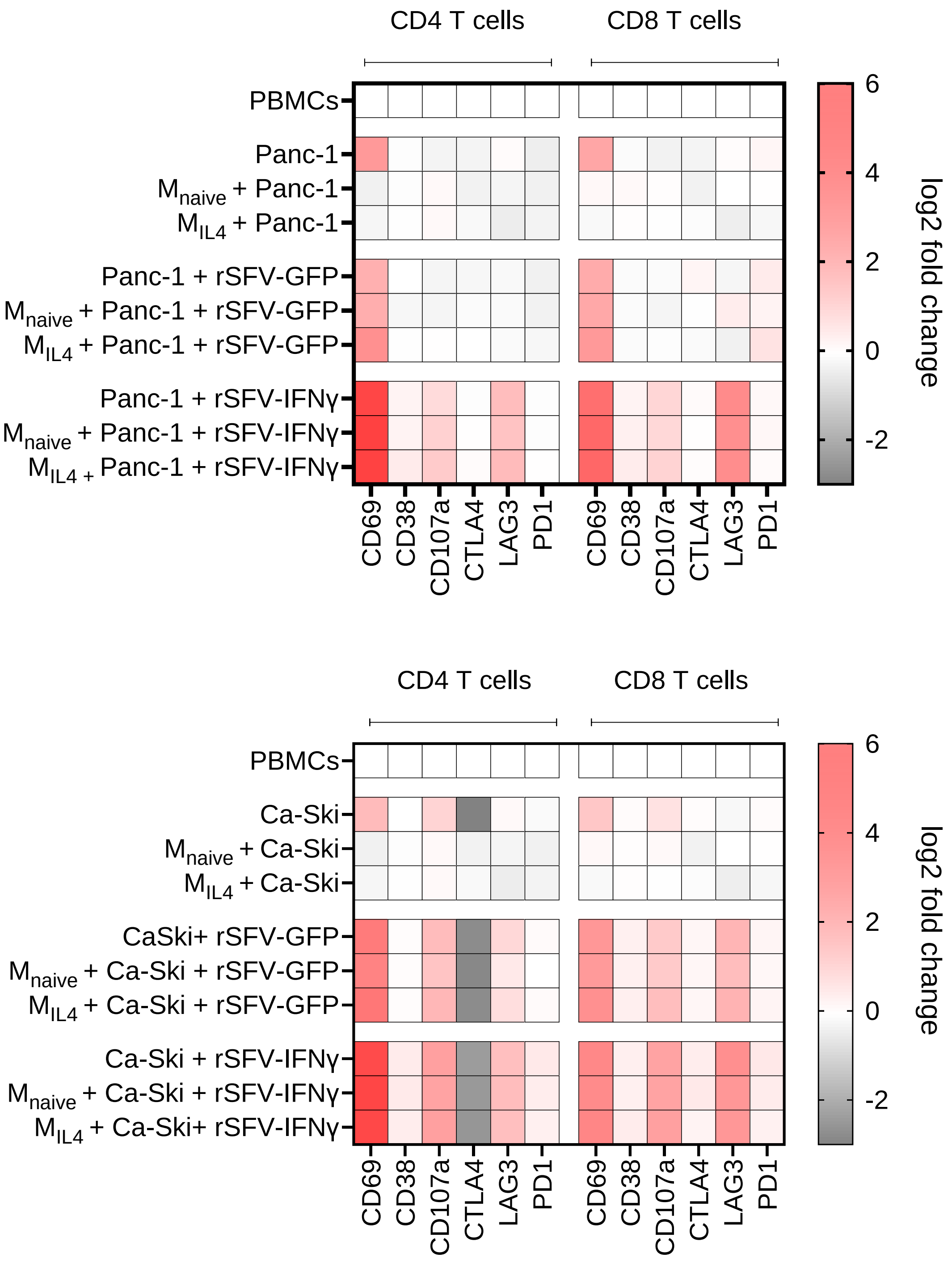


**Supplementary Figure 5:** Broad evaluation of T-cell activation and regulatory markers in tumor-immune co-cultures upon rSFV infection. The experimental setup is the same as in Figure 5.

### Supplementary Figure 6

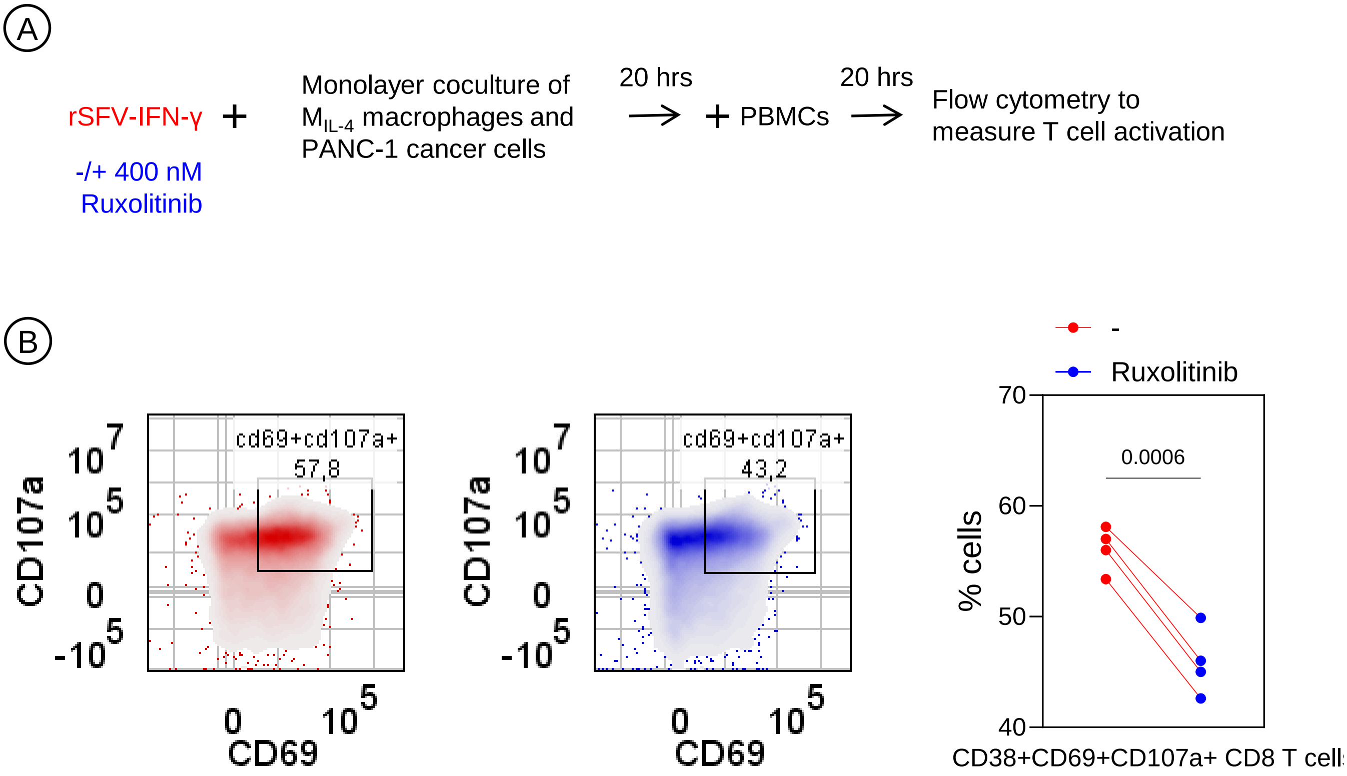


**Supplementary Figure 6:** Evaluation of T-cell activation in tumor-immune monolayer co-cultures upon rSFV-IFN-γ infection and JAK-STAT pathway blockade by Ruxolitinib. (A) Schematic representation of the experimental setup consisting of tumor-immune monolayer cocultures of cancer cells and macrophages treated with rSFV-IFN-γ, followed by the addition of PBMCs to introduce T-cells and evaluate their activation in presence or absence of 400nM Ruxolitinib. (B) Flow cytometry analysis was performed to measure activation of T-cells as explained in Figure 6. The representative flow cytometry plots on the left illustrate the quantification of CD8 cytotoxic T-cell activation (CD38+CD69+CD107a+ cells) in the tumor-immune cocultures, where rSFV-IFN-γ treated cocultures in absence of Ruxolitinib is indicated in red and in presence of Ruxolitinib in blue. Percentages represent the proportion of cells within the gate. The graph on the right represent data from 4 donors. Two-tailed paired-t-test was performed to analyze significance.

### Supplementary Figure 7

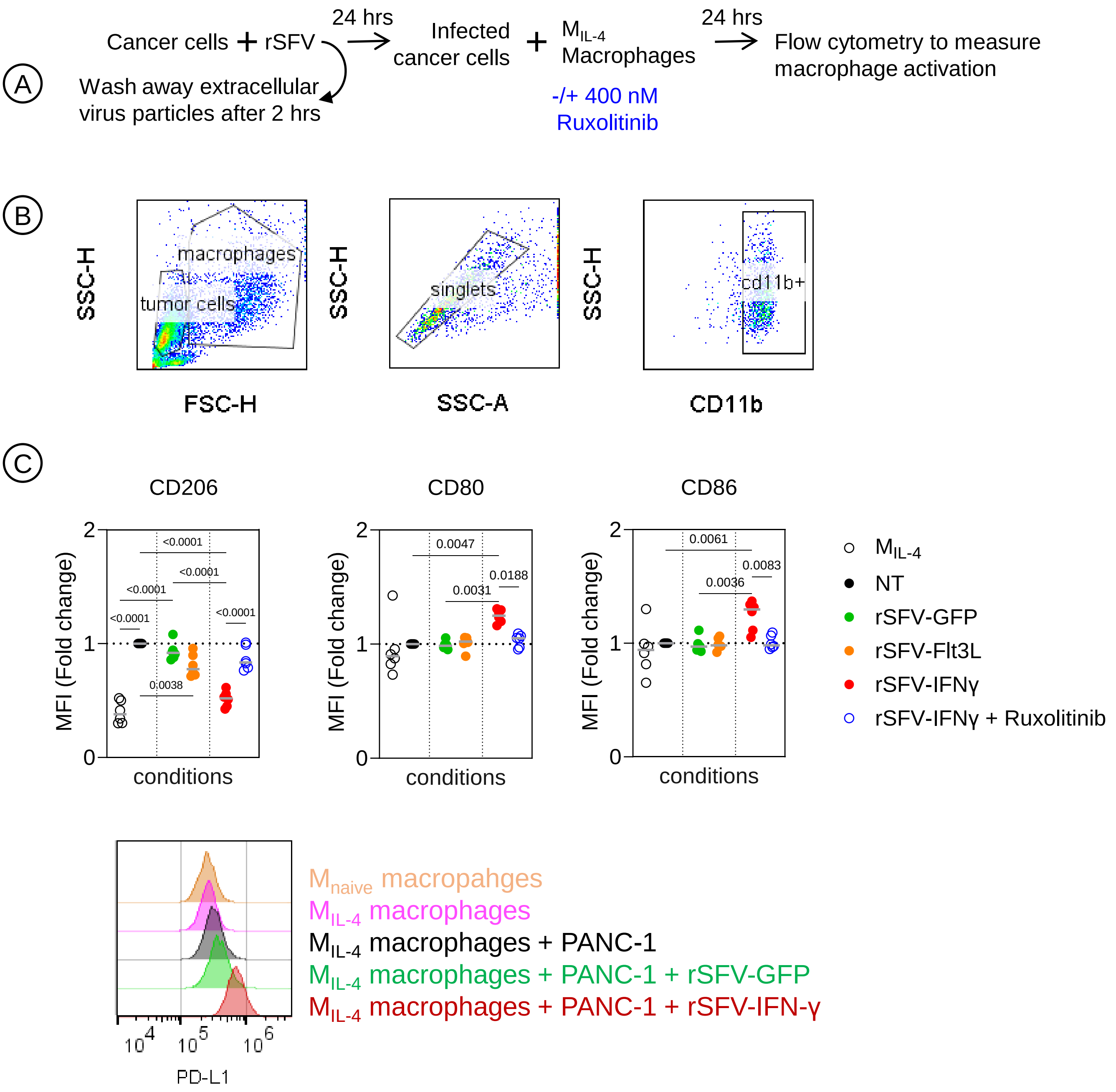


**Supplementary Figure 7:** Effect of JAK-STAT pathway on macrophage activation by rSFV-IFN-γ. (A) Experimental setup to study changes in macrophage phenotype upon stimulation by rSFV. This includes studying the effect of PANC-1 cancer cells infected by rSFV not encoding immunogenic signals (rSFV-GFP), or encoding irrelevant immunogenic signal (rSFV-Flt3L), or encoding rSFV-IFN-γ in either presence or absence of a JAK-STAT inhibitor (400nM Ruxolitinib). (B) Flow cytometry analysis of tumor-macrophage cocultures and gating strategy to study macrophage (CD11b+ cells) phenotype. (C) Change in cell-surface expression of proteins involved in either pro-tumoral (CD206, PD-L1) or anti-tumoral (CD80, CD86) function of $M_{IL4}$ macrophages cocultured or not with PANC-1 cancer cells upon stimulation resulting from rSFV-infected cells. The plots represent mean value data from 6 donors except for PD-L1 (1 donor). Significance was determined by one-way analysis of variance (ANOVA) followed by a Boneferroni post hoc test.

**Supplementary Figure 8**

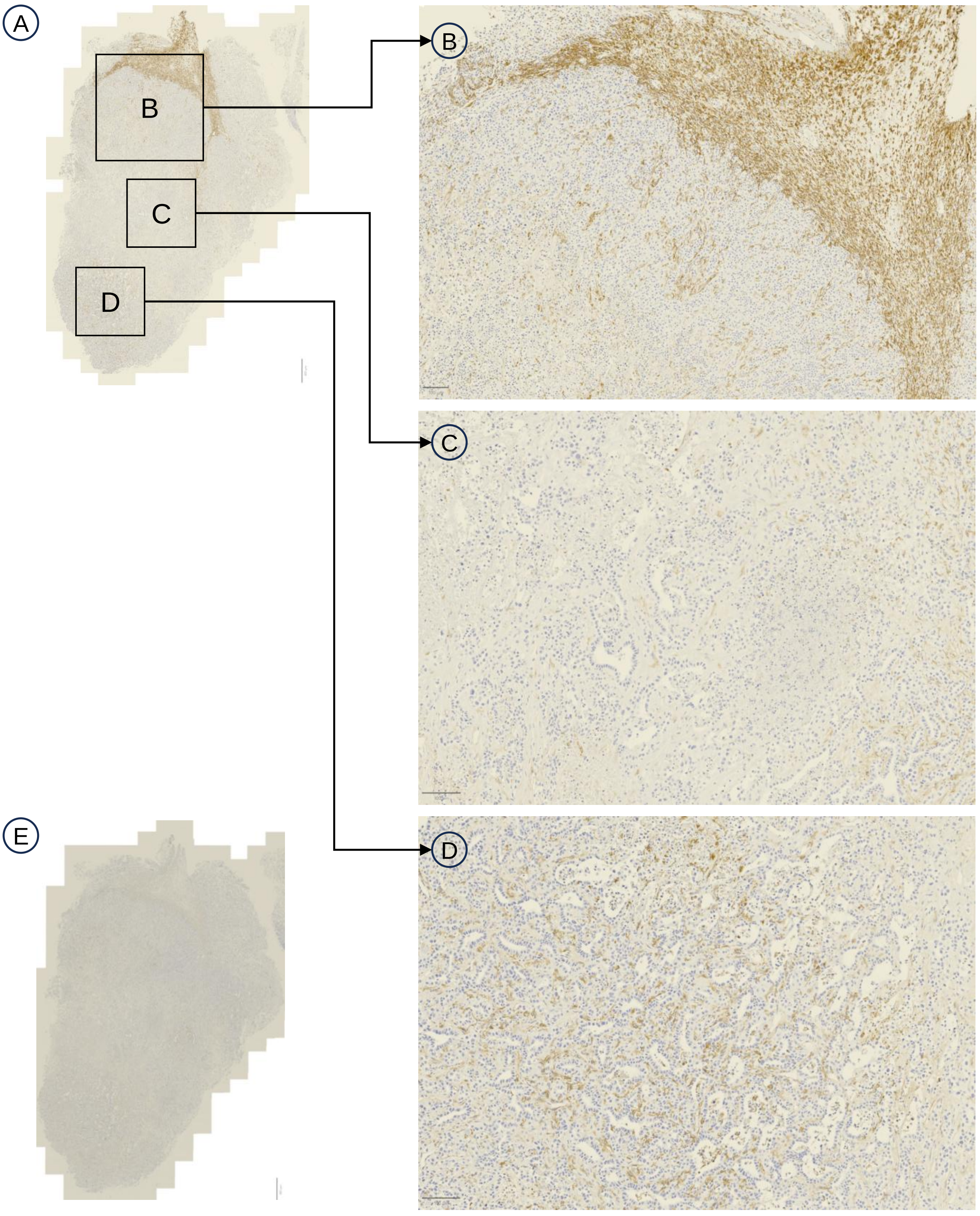


**Supplementary Figure 8:** Immunohistochemistry staining for F4/80 (brown) in the murine pancreatic cancer model KPC3. (A) Stained KPC3 tumor, 1x zoom. Squares indicate processed locations of figure S8B, C and D. (B) F4/80 expression in and around the tumor barrière, 3x zoom. (C) F4/80 expression surrounding stromal compartment, 4x zoom (D) F4/80 expression in differentiated tumor region, 4x zoom. (E) Negative control, 1x zoom. Section stained in parallel, but without primary antibody incubation.

**Supplementary Figure 9**

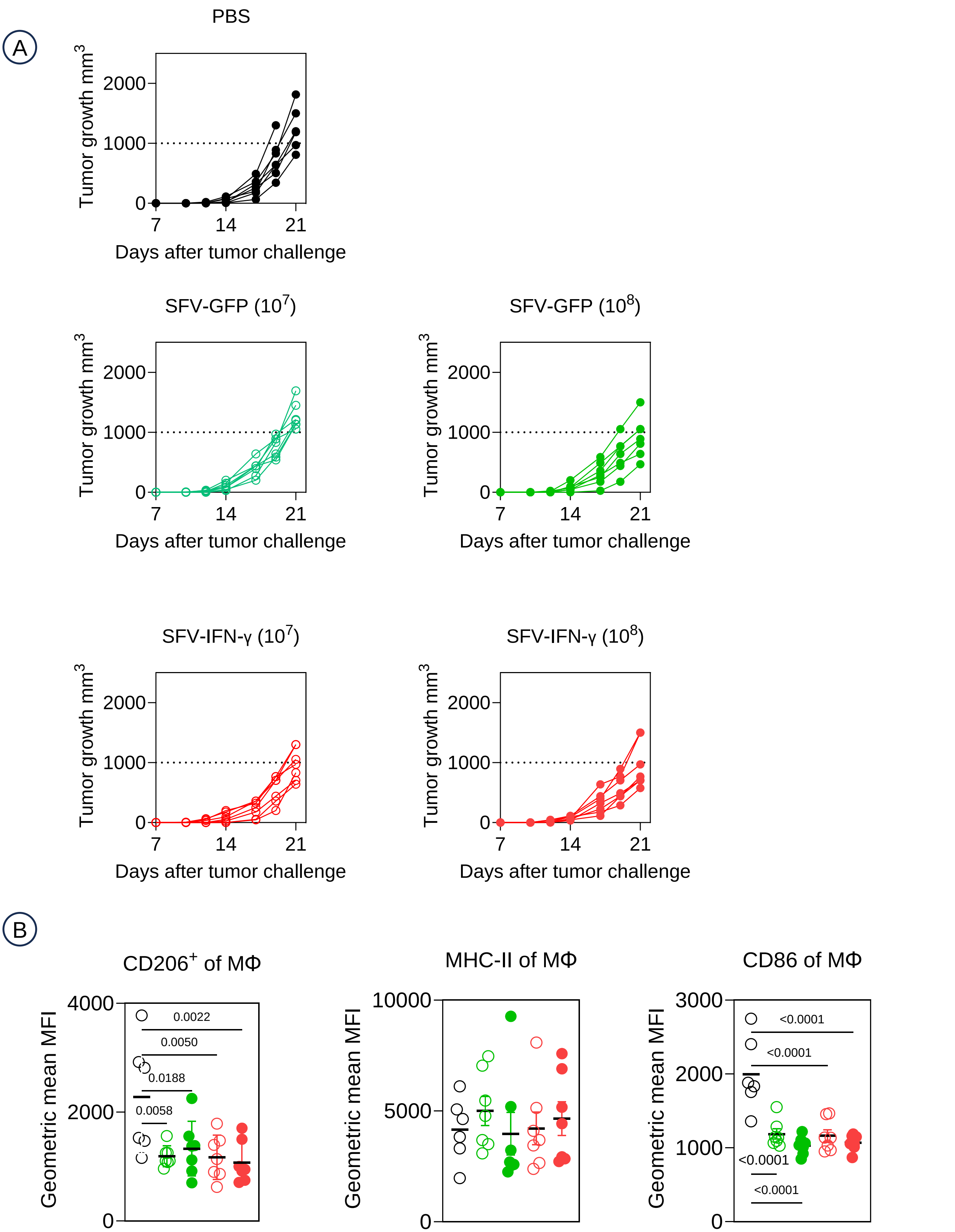


**Supplementary Figure 9: (A)** Tumor growth curves for individual animals in each treatment groups. This data corresponds to Figure 8B. **(B)** Geometric mean fluorescence based expression of CD206, MHC-II and CD86 on tumor-associated macrophages across different treatment groups.

### Supplementary Figure 10

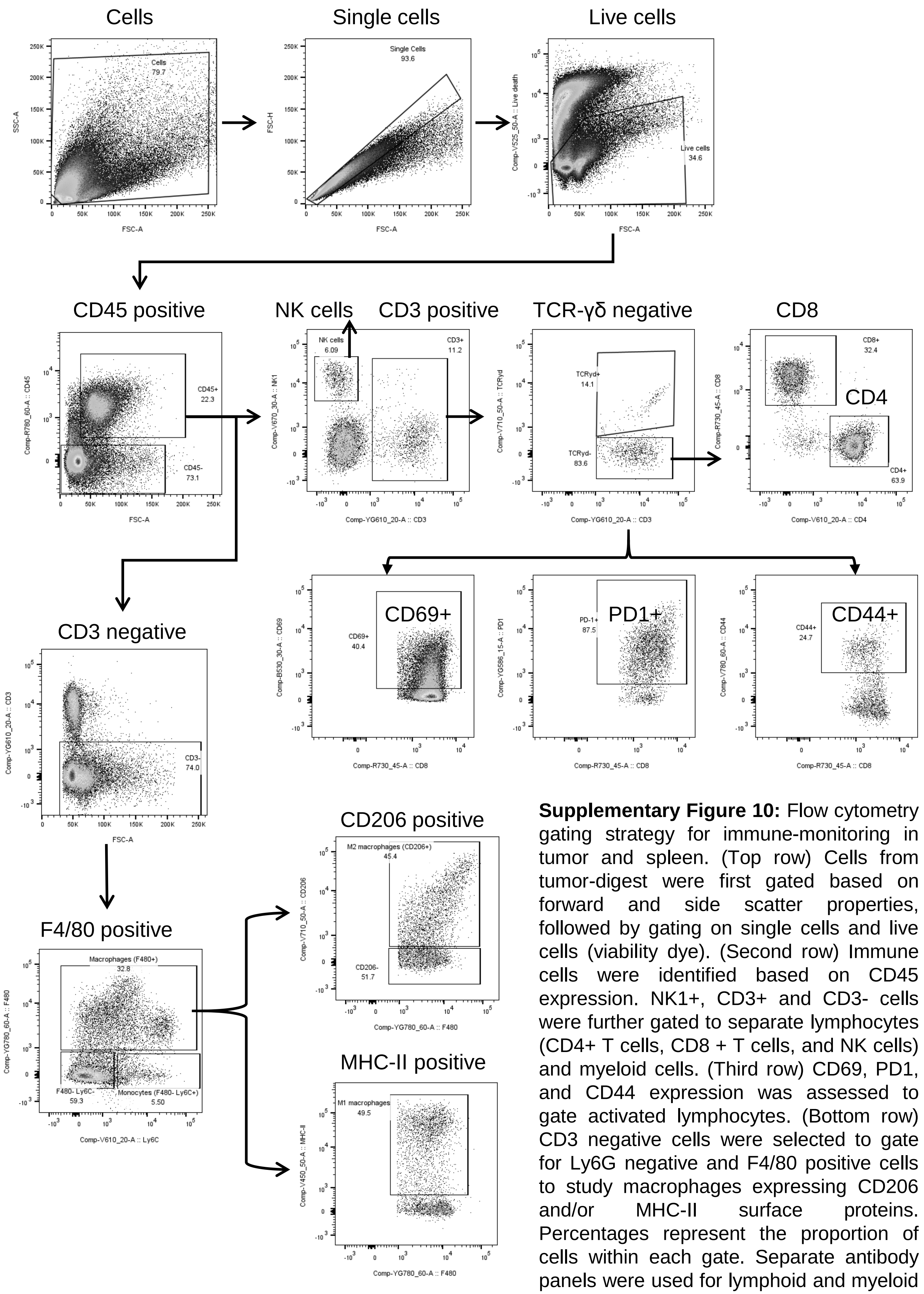


**Supplementary Figure 10:** Flow cytometry gating strategy for immune-monitoring in tumor and spleen. (Top row) Cells from tumor-digest were first gated based on forward and side scatter properties, followed by gating on single cells and live cells (viability dye). (Second row) Immune cells were identified based on CD45 expression. NK1+, CD3+ and CD3- cells were further gated to separate lymphocytes (CD4+ T cells, CD8 + T cells, and NK cells) and myeloid cells. (Third row) CD69, PD1, and CD44 expression was assessed to gate activated lymphocytes. (Bottom row) CD3 negative cells were selected to gate for Ly6G negative and F4/80 positive cells to study macrophages expressing CD206 and/or MHC-II surface proteins. Percentages represent the proportion of cells within each gate. Separate antibody panels were used for lymphoid and myeloid markers.
